# Multimolecular proofreading overcomes the activity-fidelity trade-off

**DOI:** 10.64898/2026.01.19.700236

**Authors:** Zhuo Mao, Yuanqi Jia, Yuxuan Yan, Bingze Wu, Fangzhou Xiao, Zibo Chen

**Affiliations:** State Key Laboratory of Gene Expression, School of Life Sciences, Westlake University, Hangzhou, Zhejiang, China; School of Engineering, Westlake University, Hangzhou, Zhejiang, China; Research Center for Industries of the Future, Westlake University, Hangzhou, Zhejiang, China; Institute of Basic Medical Sciences, Westlake Institute for Advanced Study, Hangzhou, Zhejiang, China; Westlake Laboratory of Life Sciences and Biomedicine, Hangzhou, Zhejiang, China

**Keywords:** kinetic proofreading, spatial proofreading, activity-fidelity trade-off, speed-accuracy trade-off, protein circuit design, multicellular circuits

## Abstract

Accurate signal processing is essential for proper cell functions, and can be achieved through kinetic proofreading, where an enzyme undergoes sequential state transitions and irreversible deactivation to enable high fidelity. However, synthetically constructing a biological proofreading system has been hindered by the difficulty in engineering single molecular state transitions. Here, we designed a protein circuit that combines diffusion and endocytosis to enable kinetic proofreading at the multimolecular level, without the conservation of total enzymes implicitly assumed in classic kinetic proofreading. Simulations revealed a previously overlooked yet experimentally crucial trade-off between circuit activity and fidelity, and theoretical analysis confirmed it to be fundamental in all kinetic proofreading systems. By integrating self-activation and mutual inhibition mechanisms, the circuit overcomes this activity-fidelity trade-off within biologically plausible parameter regimes. Our results extend proofreading schemes from single enzymes to a multimolecular context, and represent a practical and generalizable strategy for constructing high-fidelity synthetic biological circuits.

**HIGHLIGHTS:**

- We design a multimolecular and multicellular proofreading circuit
- A previously overlooked yet practically relevant trade-off arises between circuit activity and fidelity
- The activity-fidelity trade-off is fundamental in all kinetic proofreading circuits
- Self-activation and mutual inhibition mechanisms collectively overcome the activity-fidelity trade-off

## INTRODUCTION

Accurate and precise signal processing is one of the hallmarks of living systems. In the cell nucleus, DNA polymerases replicate DNA at a remarkable error rate of 10^-8^ per nucleotide.^1^ In the cytoplasm, ribosomes translate proteins at an error rate of 10^-4^ per codon.^2,3^ At the cell surface, T cell receptors recognize their cognate antigens with an error rate of 10^-6^.^4,5^ These low error rates cannot be fully explained by the free energy differences in binding cognate and noncognate substrates, raising the question of how cells carry out these processes with high fidelity.

Kinetic proofreading provides a theoretical framework for understanding the seemingly impossible high fidelity observed in many biological processes.^6–13^ Under this framework, the cognate and noncognate molecules go through discrete intermediate states before they can eventually function.^4,8,14,15^ During this process, both the cognate and noncognate molecules can be irreversibly deactivated at the expense of energy.^6,16–19^ These two mechanisms, namely time delay and irreversible deactivation, result in an exponentially amplified discriminative power based on the binding energy differences between cognate and noncognate molecules.

The high fidelity of kinetic proofreading circuits is an attractive feature of biological circuit engineering. However, most identified kinetic proofreading processes involve intricate conformational changes or modifications at the single protein level,^6,9,20–22^ which has proven challenging to design. Galstyan *et al.* proposed an intracellular spatial proofreading scheme involving molecules at the bulk level, where time delay is naturally achieved through molecular diffusion in space and irreversible deactivation is carried out by dephosphorylation.^23^ Although both the classic kinetic proofreading scheme and the proposed spatial proofreading mechanism make use of time delay and irreversible dissociation to carry out proofreading, a key difference arises: unlike classic kinetic proofreading, where time delay and irreversible deactivation happens at the single molecule level through state transition, the decoupling of time delay and irreversible dissociation happens at the multi-molecular level in spatial proofreading suggested a plausible way to build synthetic circuits (Figure S1).

Here, to maximize experimental implementability and tunability, we designed an extracellular proofreading circuit, where time delay is similarly implemented via diffusion, and irreversible dissociation is carried out by endocytosis. This circuit can proofread, but to achieve practically relevant fidelity, it suffers from low activities (defined here as protein concentrations after diffusing across a certain distance), thus revealing an activity-fidelity trade-off. By introducing new circuit architectures including self-activation and mutual inhibition, we overcame this trade-off, and identified circuits that simultaneously sustain high fidelity and activities. Additionally, our theoretical analysis revealed the activity-fidelity trade-off to be fundamental for all kinetic proofreading schemes. Our results establish a biologically plausible multi-molecular proofreading scheme, and offer design principles for proofreading circuits to overcome the activity-fidelity trade-off.

## RESULTS

### Intracellular gradient-based proofreading is difficult to establish

In spatial proofreading (Figure 1A), an enzyme binds to both cognate (R) and noncognate (W) substrates at the starting site (S_0_), giving rise to identical concentrations of both types of enzyme-substrate complexes (ER and EW). These complexes, initially inert, diffuse with the same diffusion constant through space (discretized from S_1_ to S_n-1_) towards a target site, during which the cognate and noncognate substrates dissociate from enzymes with different off rates (k_off_^ER^, k_off_^EW^). Dissociated substrates are quickly and irreversibly deactivated, after which they cannot rebind to enzymes. At the target site (S_n_), surviving ER and EW species are activated to elicit downstream signals. This process generates distinct concentration gradients for ER and EW, with their differences amplified as the distance between S_0_ and S_n_ increases.

**Figure 1.**
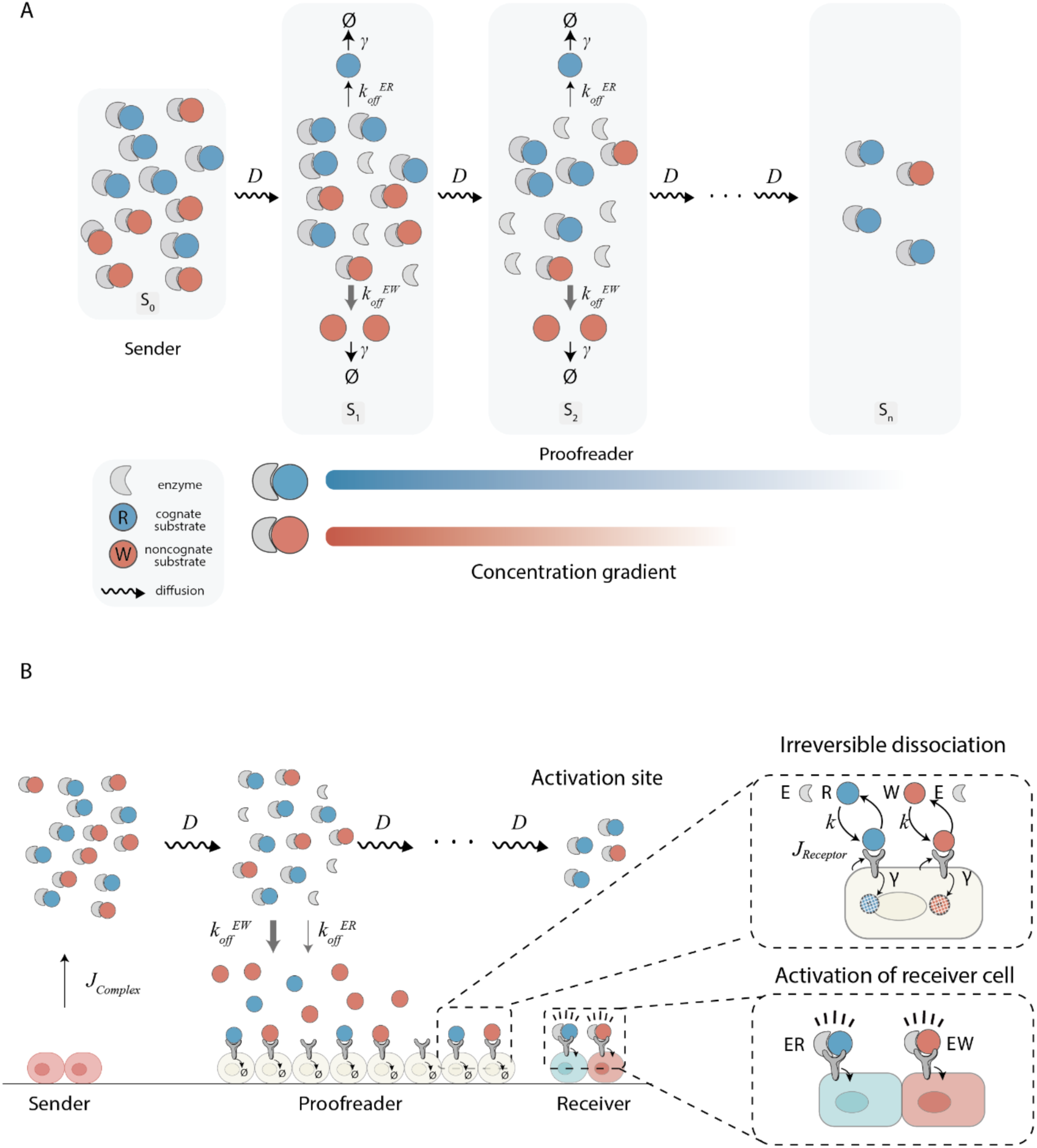
Design of the spatial proofreading system. (A) A general schematic of the spatial proofreading system. Cognate and noncognate complexes diffuse from the sender region (S_0_) to the proofreader region (S_1_-S_n-1_). Free substrates not bound by enzymes are removed at the proofreader region, generating concentration gradients. At the activation site (S_n_), located at the other end of the proofreader region, cognate and noncognate substrates are converted to their respective products. (B) Design of an extracellular proofreading circuit. Sender cells secrete cognate (ER) and noncognate complexes (EW), which undergo diffusion. Dissociated substrates are captured by endocytic receptors on the surface of proofreader cells for degradation. Both ER and EW are activated at the activation site. J_Complex_ is the secretion rate of ER and EW. D is the diffusion constant. k_off_^ER^ and k_off_^EW^ are the dissociation constants of ER and EW, respectively. R and W have the same affinity *k* with the endocytic receptor. J_Receptor_ is the synthesis rate of endocytic receptors. γ is the endocytosis rate.

The spatial proofreading scheme was originally proposed to work intracellularly.^23^ In this implementation of spatial proofreading (Figure S2A), kinases and phosphatases are spatially segregated at the membrane and cytoplasm, respectively.^24^ Cognate (R) and noncognate (W) substrates are phosphorylated by the kinases at the cell membrane, resulting in R* and W* that can bind to enzymes (E) to form cognate (ER*) and noncognate (EW*) enzyme-substrate complexes. These complexes then diffuse toward an activation site to elicit a downstream signal (e.g., transcription activation in the nucleus). As ER* and EW* diffuse in the cytoplasm, R* and W* could dissociate from the enzymes, leading to their rapid dephosphorylation back to the non-phosphorylated form by cytoplasmic phosphatases. The resulting R and W can no longer bind to enzymes. Unlike the classic kinetic proofreading model,^6,7^ where time delays are carried out via discrete molecular conformation or chemical modification changes, this scheme introduces time delay via spatial diffusion and the delayed activation of enzyme-substrate complexes.

To study the performance of the intracellular spatial proofreading scheme, we conducted simulations under varying initial concentrations of E, R, and W (Figure S2B). Here, the affinity between the cognate ER* pair is set to be 10 times higher than that of the noncognate EW* pair, comparable in magnitude to the smallest affinity difference distinguishable by classic kinetic proofreading.^6,7^ Because we are considering a bulk context where the activity comes from a collection of molecules, the fidelity performance is defined in terms of concentrations, i.e. η = [ER*] / [EW*]. In addition to fidelity, out of practical needs, we also consider activity, here defined as the concentration of ER* at the activation site. Experimentally, activity in the implementation of proofreading circuits is important, because the concentration of ER* must be high enough to be observable. Biologically, activity should also be high enough so that downstream responses are a result of proofreading, not random fluctuations or other background noises.

Simulations revealed that under a biologically plausible parameter setting (Table S1), the system maintained high fidelity at the cost of reducing the final protein concentration at the activation site to less than 0.02 nM (Figure S2B, left). This value is significantly lower than the typical binding affinity of 1 nM between a transcription factor and its target DNA binding site, hence will not elicit any downstream biological responses.^1,25^ An attempt to raise the final protein concentration by increasing the concentration of starting protein species to 10 μM resulted in a complete loss of fidelity (Figure S2B, right) with less than 1.2-fold difference between the final concentrations of ER* and EW*, despite a 10-fold difference in their binding affinities. This practical limitation provokes the question of whether there exists an alternative to achieve spatial proofreading.

### Extracellular gradient-based proofreading offers better implementability

Implementing spatial proofreading inside cells presents several practical challenges, including 1) small membrane-to-nucleus distance that makes imaging of spatial molecular gradients difficult; 2) heterogeneous intracellular environments that lead to variable diffusion constants; 3) intracellular experimental parameters (diffusion rates, diffusion distances, etc.) that are difficult to tune.^26–28^ In contrast, the extracellular environment provides an ideal platform to build and test spatial proofreading circuits by 1) offering longer distance for improved spatial and temporal resolution, 2) providing a homogeneous environment ensuring consistent diffusion, and 3) allowing key parameters to be experimentally tuned. For example, diffusion constants could be tuned by changing the extracellular media, diffusion distances can be changed by planting sender and proofreader cells at different locations. Leveraging these advantages, we propose a synthetic circuit that performs spatial proofreading in the extracellular environment.

In the extracellular proofreading model (Figure 1B), enzyme-substrate complexes (ER and EW) are secreted by sender cells at a rate of J_Complex_, and diffuse across a length L at the same diffusion rate D. Proofreader cells, expressing endocytosis receptors at a rate of J_Receptor_, intercept dissociated R and W by binding with the same affinity k, after which they are endocytosed at a rate γ, enabling irreversible dissociation (Figure 1B). Upon reaching their destination, both ER and EW molecules activate their respective downstream signals by binding to cell surface receptors. We note that all parameters here are experimentally tunable, and thus conducted numerical simulations to study the behavior of this circuit. We developed a reaction-diffusion model to describe the dynamics of this extracellular proofreading system, including substrates R (cognate) and W (noncognate), the enzyme E, their complexes (ER and EW), and membrane-bound receptor-ligand complexes (R-receptor, W-receptor). The following processes are included in our model:

1. Secretion and receptor presentation. Sender cells constitutively secrete E, R, and W into the extracellular space at zero-order secretion rates J_Complex_. The synthesis rate for the receptor is J_Receptor_.
2. Enzyme-substrate binding. Substrates R and W bind to E with an association rate constant k_on_ and dissociation rate constants k_off_ ^ER^ and k_off_ ^EW^, respectively.
3. Receptor-substrate binding. R and W undergo reversible association and dissociation with membrane-bound receptors, with kinetics governed by their respective rate constants (k_on_, k_off_^receptor^).
4. Irreversible dissociation. Proofreader cells mediate endocytosis of receptor-substrate complexes (R-receptor and W-receptor) with a first-order rate constant γ.
5. Degradation. All species (R, W, E, ER, EW, R-receptor, W-receptor, receptor) are subject to first-order degradation with a constant rate of k_deg_, representing natural turnover in the extracellular environment.

The PDEs describing this system are listed in the Methods section.

### Activity-fidelity trade-off limits the implementability of multimolecular proofreading

We first simulated a specific combination of parameters obtained from the literature.^29–31^ Similar to the intracellular proofreading scheme, the k_off_ between the cognate ER pair is set to be 10 times smaller than that of the noncognate EW pair. Additionally, the fidelity was exponentially amplified as the diffusion distance increased (Figure 2A). To provide an analytical basis for this observation, we developed a simplified model assuming that the local concentrations of the dimeric ER and EW complexes (ρ*ER*(*x*) and ρ*EW*(*x*)) are primarily governed by diffusion, dissociation, and degradation, without rebinding. Here, D denotes the diffusion coefficient of the complexes, and k_off_^R^ (or k_off_^W^) and k_deg_ represent the dissociation and degradation rates, respectively. In this way, the concentrations decay exponentially with distance x:

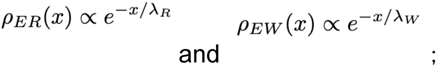

**Figure 2.**
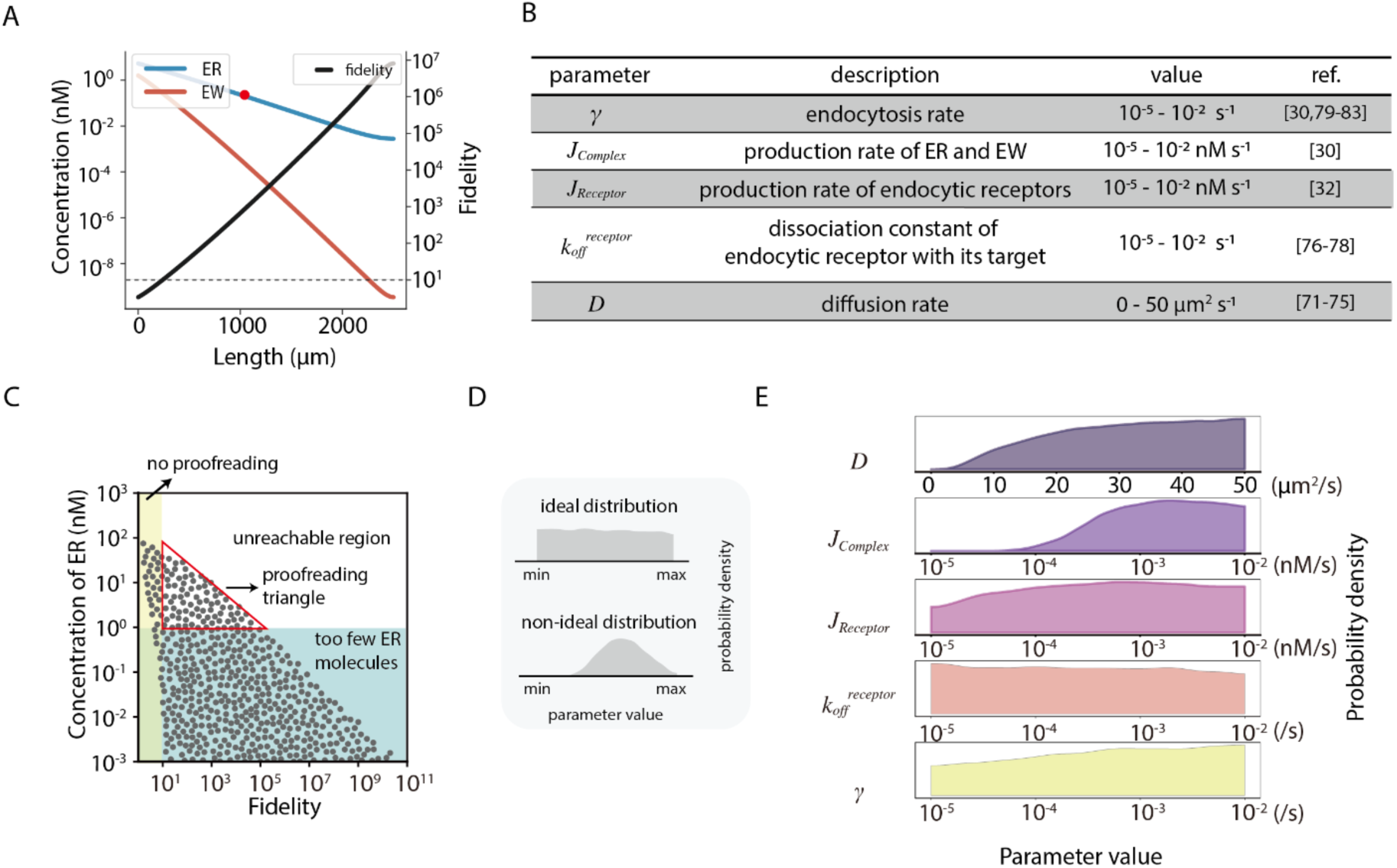
Activity-fidelity trade-off limits multimolecular proofreading. (A) Concentration gradients of ER and EW (blue and red, respectively; left axis), and the circuit fidelity of a particular parameter combination (black; right axis). Fidelity is defined as the ratio of the concentrations of ER to EW at a given location. The difference in affinities between ER and EW complexes is 10-fold, and the circuit is considered to have achieved proofreading if the fidelity exceeds 10 (dashed line). The red dot indicates the concentration of ER molecules when the fidelity is 10^3^. Parameter values used for this simulation are: J_Complex_ = 4×10^-4^ nM s^-1^, J_Receptor_= 2×10^-4^ nM s^-1^, k_off_^receptor^ = 4.5×10^-4^ nM s^-1^ k_off_^ER^ = 1×10^-4^ s^-1^, k_off_^EW^ = 4×10^-4^ s^-1^, γ = 4×10^-4^ s^-1^, and k_deg_ = 2×10^-5^ nM s^-1^. (B) Parameters and their values explored for the extracellular spatial proofreading model in this study. (C) Simulation results of the extracellular proofreading circuit. Each dot represents the fidelity and ER concentration at the activation site for a parameter combination across the range of parameter values in Figure 2B. The yellow region, where fidelity is less than 10, indicates no proofreading. The blue region, where the final concentration of ER is less than 1 nM, indicates insufficient ER to trigger downstream signaling. The gray area, representing unreachable fidelity-ER concentration combinations, highlights the activity-fidelity trade-off. The red triangle represents the proofreading regime. (D) The top panel depicts an ideal distribution of parameter values to implement a circuit, where parameters show no clear preference for certain values. The bottom panel shows a non-ideal distribution, where parameters concentrate around certain values. (E) Designability plot showing parameter distributions that give rise to simulation trajectories within the red triangle in Figure 2C. Parameter values used for this simulation are shown in Figure 2B.

where the characteristic decay lengths are given by 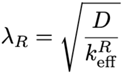 and 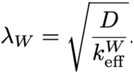, with k_eff_^R^=k ^ER^+ k_deg_ and k_eff_^W^=k_off_^EW^+ k_deg_.

Consequently, the fidelity defined as the ratio 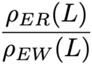 at a distance L, satisfies:

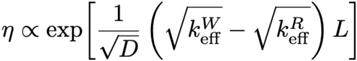

This result reveals that fidelity increases exponentially with the proofreading distance L, which is conceptually similar to how fidelity scales with the number of intermediate steps in classic kinetic proofreading schemes.^6,7^

Simulation results reveal that similar to classic kinetic proofreading schemes, the irreversible dissociation mechanism (i.e., endocytosis) is indispensable for proofreading (Figure S2C). However, achieving fidelity values above 10^3^ reduces the concentration of ER at the activation site below 0.1 nM (Figure 2A, red dot). This concentration is lower than the typical range for binding between a cell surface receptor and its ligand (nM to mM),^32–34^ making the result of proofreading incapable of eliciting any significant downstream actions. Given that both activity and fidelity need to be high enough for a meaningful proofreading behavior, a trade-off between activity and fidelity, if exists, would pose a severe limit on proofreading performance. Therefore we set off to investigate this trade-off.

To obtain a holistic view of the activity-fidelity trade-off, we performed a parameter sweep using biologically relevant values (Figure 2B). We exclude simulation results with fidelity lower than 10 (yellow area in Figure 2C), and where ER concentrations are below a typical biologically detectable threshold of 1 nM (blue area in Figure 2C). The remaining points on the plot form a “proofreading triangle”, with its upper right edge representing the Pareto-optimal front of the activity-fidelity trade-off. The area beyond the Pareto-optimal front is unreachable regardless of the choice of parameter combinations (white area in Figure 2C). This demonstrates that indeed the activity-fidelity trade-off is general for the extracellular spatial proofreading model. Additionally, simulation outcomes that fall within the proofreading triangle only constitute 28.5% of total simulation outcomes, indicating a very limited set of parameter values that could give rise to experimentally observable proofreading.

To explore the implementability of the extracellular spatial proofreading model, we analyzed parameter values that gave rise to points within the proofreading triangle. Intuitively, a wide range of parameter values in the triangle, like a uniform distribution, means significant flexibility and robustness when designing the circuit, while a narrow or peaked distribution means the contrary (Figure 2D). Parameters J_Receptor_, k_off_^receptor^, and γ exhibited nearly uniform distributions, indicating that the implementability of the circuit is robust against variations in these parameters. On the other hand, we observed a clear preference for large secretion rates of complexes and rapid diffusion.

Experimentally, the secretion rate J_Complex_ is a more tunable parameter than the diffusion constant. We therefore looked into how J_Complex_ affects the performance of the proofreading circuit. Simulation results indicate that increasing J_Complex_ resulted in expansion of the Pareto-optimal front, and enrichment in the number of parameter combinations that are capable of effective proofreading (Figure S3A). Altogether, these findings confirmed the plausibility of implementing a spatial proofreading circuit using real cells, albeit with a tight activity-fidelity trade-off. This motivated us to look for additional mechanisms to improve the spatial proofreading design.

### Self-activation could partially overcome the activity-fidelity trade-off

Having demonstrated the importance of J_Complex_ in proofreading, we explored whether modifications to the secretion rate could help overcome the activity-fidelity trade-off. We performed two simulations, where sender cells secreted ER and EW three times faster than the largest reported protein secretion rate by a mammalian cell(Figure S3B, top),^29^ or more sender cells are introduced into the system and mixed with proofreader cells, occupying the whole space under consideration (Figure S3C, top). The former scheme only gave slightly higher final ER concentrations at the proofreader site (Figure S3B, bottom), while the latter completely abolished fidelity (Figure S3C, bottom). Therefore, non-discriminatively increasing secretion could not overcome the activity-fidelity trade-off.

Given this observation, and the biological upper limit for sender cells to secrete proteins, we reasoned that proofreader cells could be engineered to bias their secretion toward the cognate complexes. Here, we note that one crucial aspect of our proofreading circuit — multicellularity — allows us to implement new mechanisms that are challenging to realize in the classic KPR scheme at the single-molecule level. We propose self-activation as a plausible strategy, where the proofreader cells sense the concentrations of diffusible cognate or noncognate molecules and respond by synthesizing and secreting detected molecules (Figure 3A). We note that this strategy also broke the implicit convention in classic KPR where the total number of enzymes stays the same, further highlighting the flexibility of the multimolecular proofreading design.

**Figure 3.**
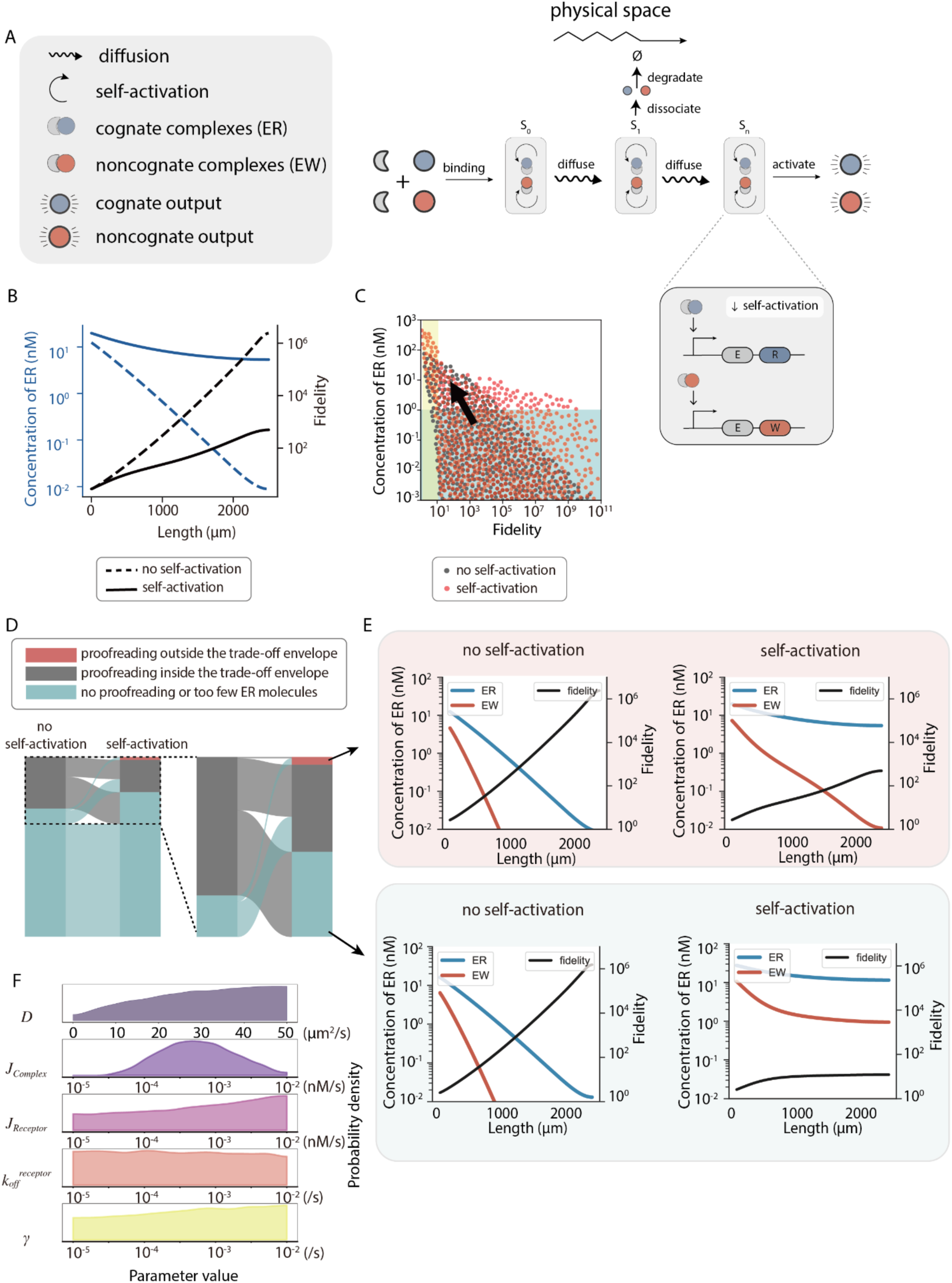
A self-activation motif partially overcomes activity-fidelity trade-off. (A) Schematic of self-activation in proofreader cells. Cognate and noncognate complexes diffuse from the sender region (S_0_) to the proofreader region (S_1_-S_n-1_). During diffusion, these complexes could dissociate into monomers, and free substrates not bound by enzymes are endocytosed at the proofreader region, generating concentration gradients. At the activation site (S_n_), located at the other end of the proofreader region, cognate and noncognate substrates are converted to their respective products. Proofreader cells possess additional receptors that sense the presence of ER and EW complexes, and respond by synthesizing and secreting ER and EW molecules, respectively. (B) Activity-fidelity plot of the proofreading circuit with self-activation. The concentration of ER increases (left axis) without diminishing the circuit fidelity (right axis). Dashed and solid lines indicate the absence and presence of self-activation, respectively. Parameter values used for this simulation are: J_Complex_ = 8×10^-4^ nM s^-1^ (equal to the maximum secretion rate in positive/negative feedback), J_Receptor_= 2×10^-4^ nM s^-1^, k_on_^receptor^ = 4.5×10^-4^ nM s^-1^, k_off_^receptor^ = 1×10^-3^ s^-1^, k_on_^ER^ = k_on_^EW^ = 1×10^-4^ nM s^-1^, k_off_^ER^ = 1×10^-4^ s^-1^, k_off_^EW^ = 1×10^-3^ s^-1^, γ = 4×10^-4^ s^-1^, and k_deg_ = 2×10^-5^ nM s^-1^. (C) Simulation results of the extracellular proofreading circuit with (red) and without (grey) self-activation. The yellow region, where fidelity is less than 10, indicates no proofreading. The blue region, where the final concentration of ER is less than 1 nM, indicates that the ER concentration is too low to elicit downstream signaling. The black arrow indicates the overall migration trajectory of simulation outcomes after adding self-activation. Parameter values for the top two groups are: J_Complex_ = 1×10^-3^ nM s^-1^ (equal to the maximum secretion rate in positive/negative feedback), J_Receptor_= 2×10^-4^ nM s^-1^, J_Receptor_= 2×10^-4^ nM s^-1^, k_on_^receptor^ = 4.5×10^-4^ nM s^-1^, k_off_^receptor^ = 1×10^-3^ s^-1^, k_on_^ER^ = k_on_^EW^ = 1×10^-4^ nM s^-1^, k_off_^ER^ = 1×10^-4^ s^-1^, k_off_^EW^ = 1×10^-3^ s^-1^, and γ = 4×10^-4^ s^-1^, k_deg_ = 2×10^-5^ s^-1^. For the bottom two are: J_Complex_ = 1×10^-3^ nM s^-1^ (equal to the maximum secretion rate in positive/negative feedback), J_Receptor_= 2×10^-4^ nM/s, k_on_^receptor^ = 4.5×10^-4^ nM s^-1^, k_off_^receptor^ = 1×10^-3^ s^-1^, k_on_^ER^ = k_on_^EW^ = 1×10^-4^ nM s^-1^, k_off_^ER^ = 1×10^-4^ s^-1^, k_off_^EW^ = 1×10^-3^ s^-1^, γ = 4×10^-4^ s^-1^, and k_deg_ = 2×10^-5^ nM s^-1^. (F) Designability plot showing parameter combinations that give rise to proofreading with sufficient ER concentrations in Figure 3C.

We model the circuit with four flavors of the self-activation arm (Figure S4A, Methods details):

1. ER and EW respectively induce the secretion of ER and EW (ER / EW → ER / EW),
2. ER and EW respectively induce the secretion of R and W (ER / EW → R / W),
3. R and W respectively induce the secretion of ER and EW (R / W → ER / EW), and
4. R and W respectively induce the secretion of R and W (R / W → R / W).

The simulation results reveal that among all four tested self-activation implementations, only the ER / EW → ER / EW scheme could increase the final concentration of ER without significantly affecting the fidelity (Figure 3B, S4B). Though, despite the introduction of self-activation, endocytosis remains essential for effective proofreading (Figure S4B), highlighting the importance of irreversible dissociation in kinetic proofreading. This scheme provides a biologically plausible way to achieve high fidelity while maintaining enough molecules for downstream signal activation.

To gain a deeper insight into how self-activation changed the landscape of activity-fidelity trade-off, we explored the entire parameter space and observed a number of simulation trajectories that overcame the previously established activity-fidelity trade-off triangle (red dots in Figure 3C). Detailed analysis of how simulation outcomes migrated in response to self-activation revealed that self-activation rescued parameter combinations that either had too few molecules or did not provide high fidelity (Figure 3D, cyan to red and black). It also drove some points from inside the proofreading triangle to the outside (Figure 3D, black to red). Interestingly, a notable number of points went from the proofreading triangle to no proofreading (Figure 3D, black to cyan), resulting in an overall decrease in the number of simulation trajectories that could achieve proofreading. Comparing with simulation trajectories that overcame the trade-off (Figure 3E, top), these points had large production rates at the sender cell region or ineffective endocytosis, which in combination with self-activation, resulted in flattened ER and EW gradients, hence low fidelity (Figure 3E, bottom).

We investigated the designability of self-activation-enabled proofreading and found that the new proofreading scheme can tolerate lower protein secretion and diffusion rates (Figure 3F). Analysis of the self-activation Hill function parameters (K, β, n, see Methods details for description) revealed that increasing these parameter values could enhance the ability of the system to overcome the activity-fidelity trade-off (Figure S5). Among these, larger β (maximal production) and n (cooperativity) increases the output and sensitivity of the Hill function, respectively; increasing the half-maximal effective concentration of the input protein (K) implicitly enhances the discrimination of cognate and noncognate complexes.

We noted that the self-activation mechanism in our model is itself a perfectly accurate signal amplifier, thus could achieve lossless boost of the proofreading activity. This may not hold in reality, since gene regulation could have error rates as well.^35^ To test whether the effect of self-activation still holds with realistic error rates, we introduced error rates in the self-activation of ER and EW. Simulation results indicate that at an error rate as low as 0.0001, the benefit of self-activation on the activity-fidelity trade-off disappeared, and the performance is worse compared to without self-activation, with fewer parameter values achieving effective proofreading (Figure S4C). This occurs because inaccurate self-activation causes a part of cognate complexes to erroneously trigger secretion of noncognate complexes at levels that match or exceed those of the cognate complexes, thereby impairing proofreading.

Taken together, by breaking the convention in classic KPR where the number of molecules stays the same and allowing proofreading cells to amplify signals by activating the expression of additional enzyme-substrate complexes, self-activation could result in KPR circuits that partially overcome the activity-fidelity trade-off. However, the advantage in performance gained is sensitive to errors in self-activation itself, therefore further improvements are needed for a design that works in practice.

### Self-activation coupled with mutual inhibition gives rise to near-perfect proofreading

Given that self-activation spatial proofreading circuits are sensitive to error, we investigated other circuit designs to further alleviate the constraints of the activity-fidelity trade-off observed in Figure 3C. The proofreading circuit with self-activation was prone to error because ER could cause significant expression of EW by mistake. To counter this, we aimed to develop strategies ideally allowing ER to inhibit the expression of EW, therefore compensating the erroneous activation effect. We first considered a mutual inhibition scheme, where proofreader cells secrete ER and EW at the same basal rate in the absence of inputs but can be inhibited by EW and ER, respectively (Figure S4D). Because of the diminishing concentration of ER and EW at the final activation site, their inhibitory effect on the proofreader cells becomes minimal, resulting in almost the same level of secretion of EW and ER, hence very low fidelity (Figure S4E).

Inspired by winner-take-all circuits,^36–40^ we combined self-activation and mutual inhibition (SAMI), in which ER / EW could self-activate and mutually inhibit in the proofreader region (Figure 4A). Notably, when simulating across the entire parameter space, SAMI results in almost perfect proofreading, where the activity-fidelity trade-off was nearly completely eliminated (Figure 4B). Analysis of point migration revealed a bigger percentage of points (44.5%) that overcame the trade-off than that from self-activation alone (19.5%) (Figure 4C), and improvements in both the final concentration of ER and the overall fidelity (Figure 4D) for a wide range of parameter combinations (Figure 4E). We also investigated the designability of SAMI-enabled proofreading and found that the new proofreading scheme can tolerate a broader range of protein secretion and diffusion rates than the SA-enabled proofreading (Figure 4E).

**Figure 4.**
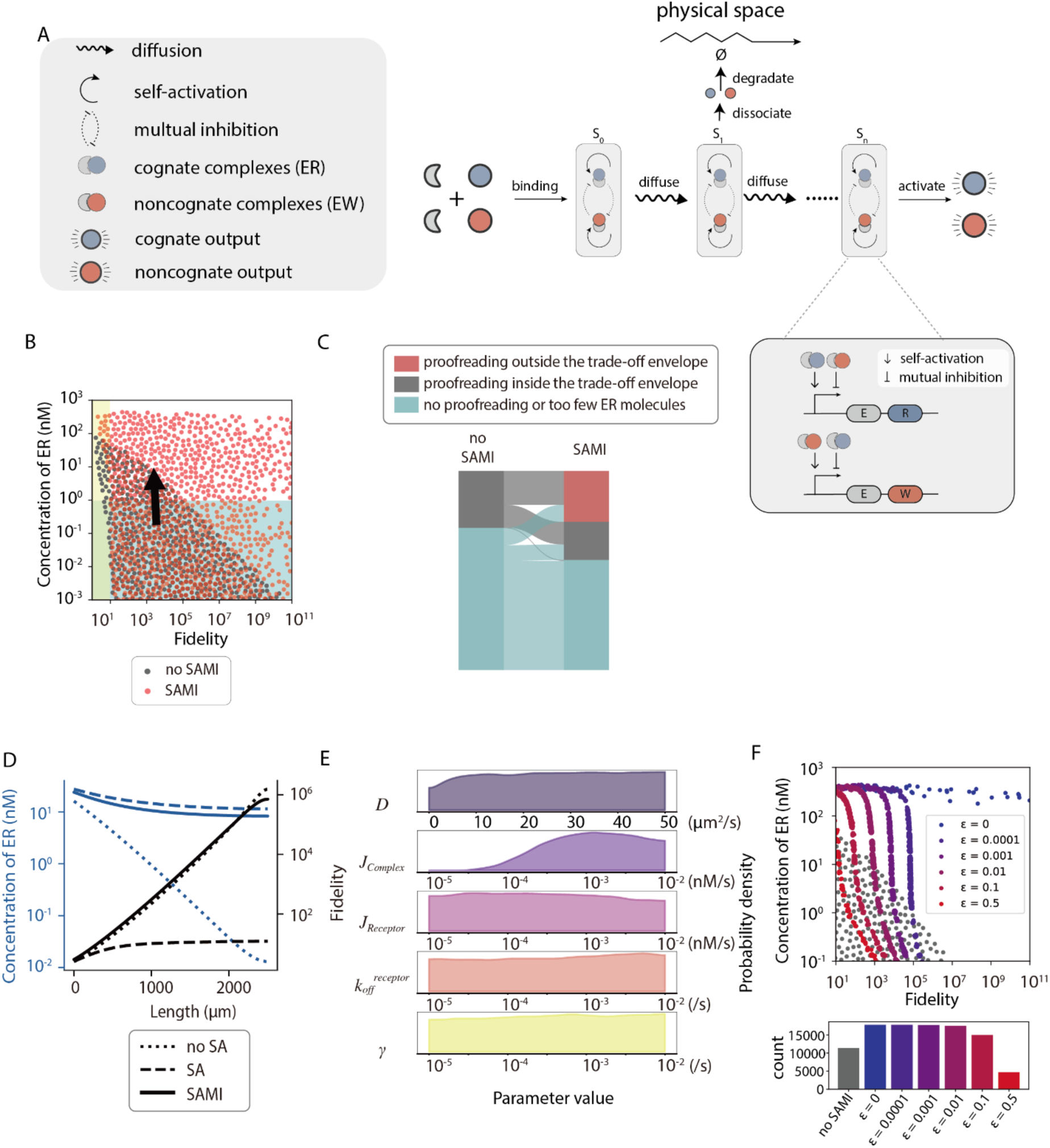
Self-activation with mutual inhibition circuit eliminates the activity-fidelity trade-off. (A) Schematic of self-activation with mutual inhibition (SAMI) in proofreader cells.Cognate and noncognate complexes diffuse from the sender region (S_0_) to the proofreader region (S_1_-S_n-1_). During diffusion, these complexes could dissociate into monomers, and free substrates not bound by enzymes are endocytosed at the proofreader region, generating concentration gradients. At the activation site (S_n_), located at the other end of the proofreader region, cognate and noncognate substrates are converted to their respective products. Proofreader cells possess additional receptors that sense the presence of ER and EW complexes and respond by synthesizing and secreting ER and EW molecules, respectively. Meanwhile, ER and EW can inhibit the synthesis and secretion of each other. (B) Simulation results of the extracellular proofreading circuit with (red) and without (gray) SAMI. The yellow region, where fidelity is less than 10, indicates no proofreading. The blue region, where the final concentration of ER is less than 1 nM, indicates that the ER concentration is too low to elicit downstream signaling. The black arrow indicates the overall migration trajectory of simulation outcomes after adding SAMI. (C) River plot indicating the migration of simulation outcomes when SAMI was introduced to the circuit. Gray region indicates points that fall within the activity-fidelity trade-off envelope. Blue region indicates too low of a fidelity or ER concentration. Red region indicates the newly emerged regime where simulation results have overcome the activity-fidelity trade-off. (D) Comparison of proofreading circuits with no regulation, with self-activation, and with SAMI. Dotted, dashed and solid lines indicate no regulation, with self-activation, and with SAMI, respectively. Parameter values used for this simulation are: J_Complex_ = 1×10^-3^ nM s^-1^ (equal to the maximum secretion rate in positive/negative feedback), J_Receptor_= 2×10^-4^ nM s^-1^, k_on_^receptor^ = 4.5×10^-4^ nM s^-1^, k_off_^receptor^ = 1×10^-3^ s^-1^, k_on_^ER^ = k_on_^EW^ = 1×10^-4^ nM s^-1^, k_off_^ER^ = 1×10^-4^ s^-1^, k_off_^EW^ = 1×10^-3^ s^-1^, γ = 4×10^-4^ s^-1^, and k_deg_ = 2×10^-5^ nM s^-1^. (E) Designability plot showing parameter combinations that give rise to proofreading with sufficient ER concentrations in Figure 4B. (F) Activity-fidelity trade-off for the SAMI circuit across various error rates. The scatter plot shows the activity-fidelity trade-off boundary for a range of error rates (ε) in the SAMI circuit: ε = 0, ε = 0.0001, ε = 0.001, ε = 0.01, ε = 0.1. The bar plot shows the count of simulation trajectories capable of proofreading in each error rate category.

To investigate the SAMI circuit in a more biologically realistic setting, we incorporated error rates in both self-activation and mutual inhibition of ER and EW. Distinct from the self-activation circuit, where only a slight error could destroy the newly gained improvement in the activity-fidelity trade-off (Figure S4C), error rates only affected the maximal fidelity without decreasing the overall activities of SAMI circuits (Figure 4F). Additionally, even at 10% error rate (ε = 0.1), SAMI still produced more simulation outcomes with both high proofreading abilities and elevated ER concentrations, compared to the proofreading circuit without SAMI (Figure 4F, bottom). These findings underscore the robustness of SAMI circuits under error-prone conditions.

We performed additional simulations to investigate the effect of differential secretion rates by sender cells. When EW is expressed at rates ranging from 32 to 256 times higher than that of ER, the system initially suffers from low fidelity (Figure S7, left). However, this trend could be reversed with increasing diffusion distance, thanks to the error-correcting mechanism offered by endocytosis. Additionally, the introduction of SA decreased the distance needed for this error correction (Figure S7, middle), while SAMI additionally enhanced the overall fidelity (Figure S7, right). These results underscore the versatility and robustness of the circuit across a wide range of secretion imbalances.

We investigate the time, material, and energy costs associated with SAMI. Panels A and B in Figure S8 respectively show the dynamics of ER concentration and fidelity at a distance L = 2000 µm, for spatial proofreading circuits with and without SAMI. Although both systems reached the 1,000 threshold for fidelity at similar times (Figure S8B), SAMI reached the target concentration of 0.1 nM much faster than the circuit without SAMI (Figure S8A). Therefore, the addition of SAMI reduces the overall time cost of the proofreading system. We define material cost as the total production of E, R, and W required for the spatial proofreading system to reach the activity and fidelity thresholds in Figure S8A and B. Figure S8C shows the cumulative production of E, R and W as a function of time for the circuit with and without SAMI, taking into consideration basal production by sender cells and dynamically changing production rates by proofreader cells due to SAMI. Since the circuit with SAMI reaches the activity and fidelity thresholds much faster (Figure S8A, B), it also resulted in a lower cumulative production of proteins. Hence, under this parameter regime, SAMI also reduces the material cost. However, energy consumption, in particular when live cells are involved, is difficult to define precisely. The additional processes, such as endocytosis^41–43^ and transcriptional regulations^44^, could add energetic burdens to cells.

Collectively, the multimolecular nature of our design allowed the flexible introduction of additional layers of regulation, resulting in elimination of the activity-fidelity trade-off. Given the importance of the multimolecular nature of our design to overcome the activity-fidelity trade-off, we wondered whether the multimolecular design itself is a general and necessary strategy to implement proofreading schemes in practice. This constitutes an investigation of activity-fidelity trade-offs in traditional proofreading schemes, which we describe in the next section.

### Activity-fidelity trade-off is fundamental in kinetic proofreading

An ideal proofreading circuit should process biological signals with high fidelity and speed, while expending minimal energy. It should also produce outputs with enough abundance or activity to elicit downstream signals. In most studies, fidelity was defined as the ratio of cognate to noncognate product concentrations,^6,7^ or the probability of correct versus incorrect outcomes;^8,45^ speed was represented by the first-passage time of cognate complexes;^16,17,45^ energy was defined as the free energy required to drive the proofreading process through intermediate states, which scales with the number of intermediate states (Table S2). Extensive theoretical and experimental studies on the relationship among these three performance metrics have revealed that improving one metric is often done at the cost of deteriorating either or both of the other two metrics.^6,7,16,45–50^ However, activity, which was shown above as a practically important performance metric, has largely been overlooked.

Our numerical simulations have identified activity-fidelity as an indispensable trade-off for multi-molecular proofreading circuits. In order to explore whether this additional trade-off also applies to single-molecular proofreading systems, we analyzed a generalized KPR circuit involving enzymes and their cognate and noncognate substrates (Figure 5A).^6,8,15^ In this model, key parameters include dissociation rates for cognate (k_off_^ER^) and noncognate (k_off_^EW^) substrate-enzyme binding, the association rate between substrates and enzymes (k_on_), the transition rate between intermediate states (k_transit_), and the catalytic rate for product formation (k_cat_). To encompass variations of KPR circuits involving different final product activation mechanisms,^8,9,14,51–56^ we define activity as the concentration of cognate complexes at the final intermediate state (ER_n_), right before they are converted into products by enzymes, or elicit downstream signaling as full complexes. Fidelity is defined as η_n_ = [ER_n_] / [EW_n_], the ratio of the concentrations of cognate (ER_n_) and noncognate (EW_n_) complexes at the final intermediate state. Our model assumes that the total concentration of enzymes (free enzymes plus enzyme-substrate complexes) is a constant c_0_. We use c’_n_ in the following calculation as the normalized concentration, i.e. c’_n_= c_n_ / c_0_. For an n-step proofreading circuit, final activity (c_n_) and fidelity (η_n_) are expressed as (see Supplemental Text S1 for detailed derivations):

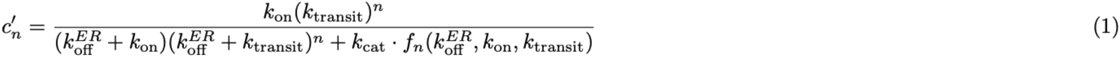

**Figure 5.**
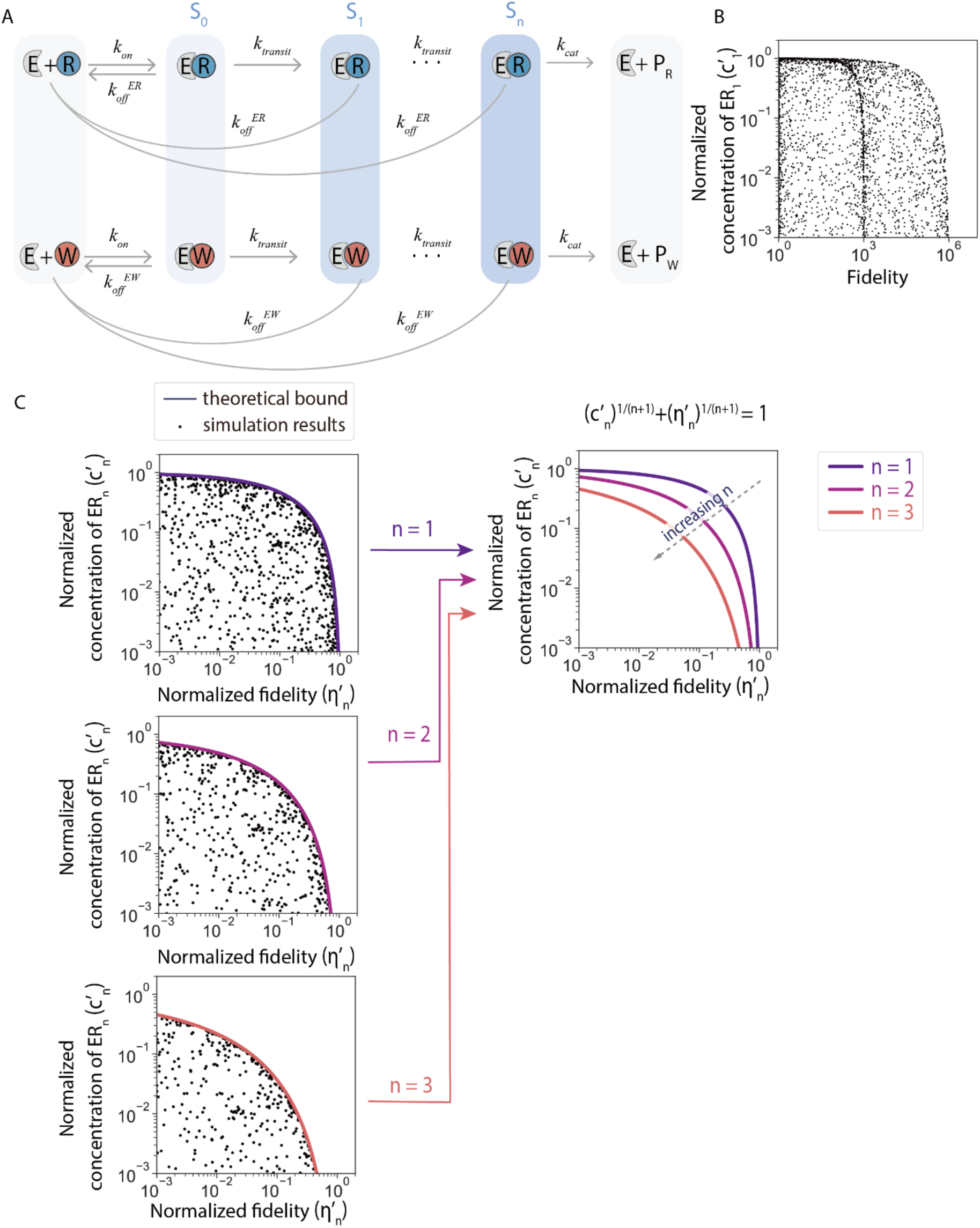
Activity-fidelity trade-off is fundamental in kinetic proofreading. (A) A schematic of classic single molecular KPR. Cognate (blue) and noncognate (red) substrates at the initial state (S_0_) bind to enzymes with an association rate of k_on_ and dissociation rates of k_off_^ER^ and k_off_^EW^, respectively. Bound complexes undergo sequential state transitions at a rate of k_transit_. At each intermediate step (S_1_-S_n_), irreversible dissociations may occur. At the final intermediate state (S_n_), cognate and noncognate substrates are catalytically converted to their respective products (P_R_ and P_W_) at a rate of k_cat_. (B) Activity-fidelity trade-off of the KPR circuit with one intermediate state (n = 1). Black dots represent simulation outcomes with various parameter value combinations. The Y-axis shows the normalized concentration of ER (c’_n_), which is calculated as the concentration of the ER complex at step n divided by the total concentration of enzymes (free enzymes plus enzyme-substrate complexes). The X-axis shows the normalized fidelity (η’_n_), calculated as the overall fidelity η_n_ normalized by the maximal fidelity achievable with an n-step KPR (σ^n+1^), where σ = k_off_^EW^ / k_off_^ER^ = 1,000 is the ratio of the dissociation rates between EW and ER. All parameters used and their values are shown in Table S3. (C) Analytical solutions for the trade-off boundaries between the normalized activity (c’_n_) and normalized fidelity (η’_n_). Black dots represent simulation outcomes with various parameter value combinations. The solid lines are theoretical activity-fidelity trade-off boundaries for n = 1, 2, 3. The superposition of all three theoretical trade-off boundaries are shown on the right. Parameter values used for this simulation are shown in Table S1.

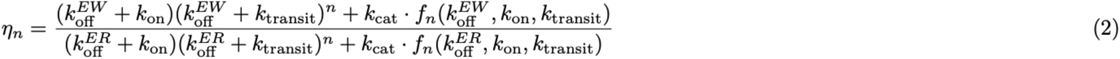

Here, f_n_ refers to:

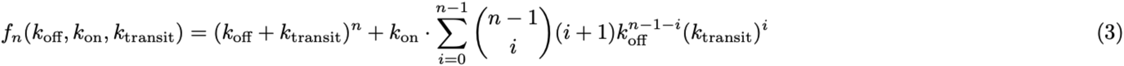

Assuming enzymes are saturated by substrates, we conducted a parameter sweep for the simplest KPR circuit with only one intermediate state (n = 1). Here, we assumed no competition between the cognate (R) and noncognate (W) substrates for enzyme binding. A clear trade-off emerged between the concentration of ER and circuit fidelity, which persisted regardless of the choice of parameter values (Figure 5B). When taking into account the competition between R and W for enzyme binding, simulation results similarly revealed a clear trade-off boundary (Figure S9A, see Supplemental Text S2 for detailed derivations). In order to precisely define this trade-off boundary, we employed a constrained optimization framework and derived an analytical expression for the trade-off curve (Figure 5C, see Supplemental Text S1 for full derivations):

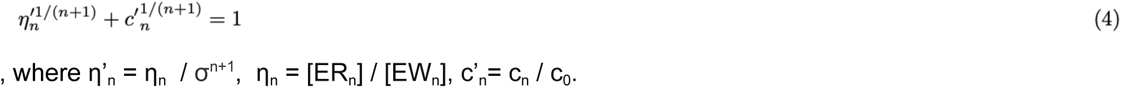

Here, η’_n_ is the overall fidelity η_n_ normalized by the maximal fidelity achievable with an n-step KPR (σ^n+1^). σ is the ratio of the dissociation rates between EW and ER, which is the fidelity at thermodynamic equilibrium. The trade-off between activity and normalized fidelity persists as n increases (Figure 5C, right). This analytical expression of the activity-fidelity trade-off can be broadly applied across different implementations of kinetic proofreading, thus establishing the activity-fidelity trade-off as a universal feature of KPR. This trade-off is also fundamental, as it could not be overcome by modifying parameter values (Figure 5C).

These results, combined with our similar findings when designing multimolecular KPR circuits (Figures 2-4), confirms the activity-fidelity trade-off as a fundamental trade-off in all KPR systems. Taken together, our theoretical analysis emphasizes the generality and necessity of multimolecular KPR circuits, where one could break the constraint on the conservation of total molecules by introducing additional regulatory layers such as SAMI, thus overcoming the activity-fidelity trade-off.

### Spatial proofreading in natural morphogenic systems

Some natural morphogens may utilize similar principles to regulate their gradients. For example, mechanisms similar to the spatial proofreading scheme shown in this paper could play a role in the regulation of BMP gradients in *Drosophila* development. In the BMP pathway, BMP-4 and BMP-7 proteins show slightly different binding affinities toward the Chordin protein, which binds BMP and protects them from degradation during diffusion.^57–59^ Our simulations show that, akin to our extracellular spatial proofreading scheme, the presence of Chrodin could reshape the gradients of BMP-4 and BMP-7, resulting in an exponentially increasing difference in their concentrations as the diffusion distance increases (Figure S10). Additionally, negative and positive feedback mechanisms, observed in morphogen gradient systems like the formation of long-range Nodal and Lefty gradients in mouse embryos,^60^ where Nodal and Lefty induce their own expressions and Lefty inhibits the expression of Nodal, hint at SAMI-like mechanisms in morphogen gradient formation.

## DISCUSSION

Intrigued by the single molecular KPR scheme,^3,6,7^ we sought to determine whether similar principles could be harnessed to design KPR at the circuit level. Building on the spatial proofreading framework (Figure 1A),^23^ we conceptualized a system where intracellular proofreading principles are extended to the multicellular level, leveraging time delays imposed by diffusion and irreversible dissociation via endocytosis. While this approach could give rise to high fidelity, simulations revealed a critical constraint: rapid depletion of molecules resulted in their low concentrations at the final activation site, oftentimes unable to elicit physiological responses (Figure 2A & 2C).

Our theoretical analysis on the activity-fidelity trade-off showed that this activity-fidelity trade-off is universal, applicable to both single- and multi-molecular KPR circuits; it is also fundamental, meaning it could not be overcome simply by tuning parameter values (Figure 5C). This provokes the question of whether changes in circuit architecture could help overcome this practically relevant trade-off.

We incorporated the SAMI module into our circuit, which uses self-activation to compensate for the depletion of molecules, and mutual inhibition to amplify the differences between cognate and noncognate complexes (Figure 4A). The kinetic proofreading enhanced by SAMI gave rise to nearly perfect proofreading with minimal trade-off between activity and fidelity (Figure 4B).

In T-cell receptor signaling, prior studies have explored mechanisms to enhance signal amplification while preserving selectivity, which bear some similarities to the activity-fidelity trade-off we investigate here. For instance, McKeithan^8^ introduced the kinetic proofreading model in T-cell receptor activation, highlighting that high activity and fidelity could not be simultaneously achieved due to inherent biochemical constraints. Subsequent studies, such as those from Altan-Bonnet and Germain,^61^ Deshek et al.,^62^ Francois et al.,^63^ and White et al.,^56^ built upon this foundation by identifying mechanisms that amplify weak signals while maintaining selectivity in T cell signaling. Specifically, Francois et al.,^63^ proposed a model that improves selectivity, where the phosphorylated TCR complexes activate SHP-1, which dephosphorylates components of the TCR complex, generating a negative feedback. These studies have indeed inspired us to use negative and positive regulations to overcome the fidelity-activity trade-off in our synthetic circuit (Figures 3 and 4).

However, there is a key difference between our work and previous studies: we distinguish between chemical state transitions (as seen in classic kinetic proofreading) and spatial state transitions (Figure S1). As the T-cell activation mechanism in these studies relies on negative or positive feedback loops by complex biochemistry, such as phosphorylation or allostery, it poses significant challenges for experimental implementation. Therefore, we chose to focus on spatial proofreading as a practical approach for synthetic circuit design.

Our circuit design offers three advantages. First, its multimolecular nature allows the flexible introduction of regulations such as self-activation and mutual inhibition mechanisms. This is in contrast to single molecular proofreading,^5,8,16,17,64,65^ where similar designs could be difficult to deploy at the single-molecule level. Second, the nature of the multicellular circuit allows energy to be spent on self-activation and mutual inhibition, in addition to driving the proofreading process through intermediate states as seen in single molecular KPR. This helps to preserve circuit activities and fidelity, leading to the elimination of the activity-fidelity trade-off. Lastly, the core of the proofreading mechanism, namely time delay and irreversible dissociation, acts indiscriminately on both cognate and noncognate complexes. Therefore, the circuit makes no assumptions about right and wrong substrates. This allows the same circuit design to be generalized to distinguish between various combinations of substrates as long as there are differences in their binding kinetics.

The incorporation of SAMI entails increased energy consumption, suggesting that future studies should examine additional trade-offs when energy constraints are factored in. Furthermore, our models did not account for biological noise, whether intrinsic or extrinsic. Investigating the performance of the circuit and its variants under conditions of low molecule counts would therefore be another avenue for future research.

In terms of experimental implementation, our design uses mechanisms commonly found in synthetic protein circuits,^66^ such as binding, dissociation, and endocytosis. This allows the circuit to be built without requiring intricate single-molecular structural designs. Importantly, the designability analysis of our circuit provides experimentally testable guidelines for tuning secretion rates, diffusion rates, receptor densities, and endocytosis rates (Figure 2E, 3F, 4E). Our design therefore provides a flexible, versatile, and generalizable framework for developing high-fidelity biological circuits.

## STAR★Methods

### Key resources table

| REAGENT or RESOURCE | SOURCE | IDENTIFIER |
| --- | --- | --- |
| **Deposited data** |  |  |
| Simulation parameters and results | This paper | https://datahub.wllsb.edu.cn/#/datade tail/465a1f160d6a4ae6b08800bae493b954 |
| **Software and algorithms** |  |  |
| Source code for ODE modeling, simulation and visualization | This paper | https://github.com/BioProgramming-Lab/proofreading-2024-12 |
| Python v3.12.2 | Python Software Foundation | https://www.python.org/ |
| NumPy v1.26.4 | NumPy developers | https://numpy.org/ |
| SciPy v1.15.3 | SciPy developers | https://scipy.org/ |
| Matplotlib v3.9.2 | Matplotlib developers | https://matplotlib.org/ |
| Zarr-Python v3.0.3 | Zarr developers | https://zarr.dev/ |

## Methods details

### Construction of an intracellular spatial proofreading model for kinases-phosphatases reaction

To investigate the dynamics of intracellular phosphorylation and dephosphorylation, we adapted the spatial proofreading model of Galstyan *et al*.^23^ This model aimed to construct an intracellular spatial proofreading model based on protein-protein interaction, protein diffusion, and phosphorylation events.

#### Model description and assumption

In this model, membrane-bound kinases phosphorylate intracellular substrates R (cognate) and W (noncognate) at the same rate (k_kin_) on the membrane, changing R and W into active forms R* and W*. R* and W* subsequently bind to enzymes E with the same on rate (k_on_) to form ER* and EW*. These complexes are assumed to diffuse uniformly throughout the whole intracellular space with a constant diffusion coefficient (D). Meanwhile, free R* and W* can be dephosphorylated by phosphatases (k_p_) in the cytoplasm when dissociated from complexes with their respective off rates (k_off_^ER^ and k_off_^EW^) (Figure S2A). The fidelity (η) is defined as the ratio of correctly formed complexes (ER*) to incorrectly formed complexes (EW*) at the activation site. We consider a mass-conserving system in which the change in total concentrations of cognate ([R] + [R*] + [ER*]) and noncognate ([W] + [W*] + [EW*]) molecules equals zero with no synthesis and degradation. Given the spatial localization of kinases at the membrane, we define the boundary conditions:

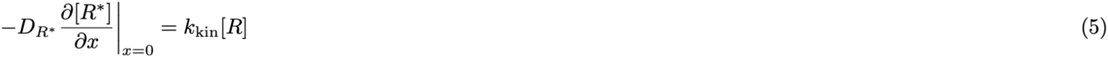

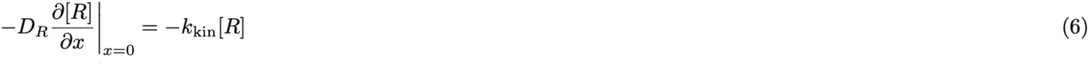

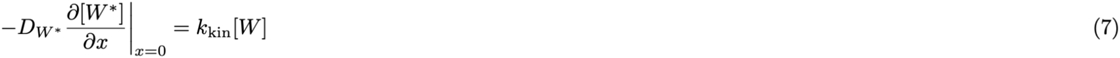

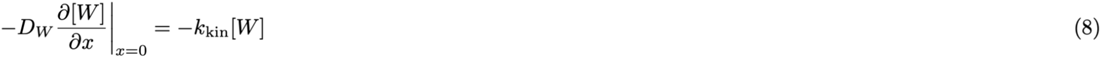

We consider zero flux for other species at the boundary. Assuming the same diffusion constant for all protein species, we characterize the reaction-diffusion system in one dimension. The diffusion term is approximated by the expression D∇^2^C, where C denotes the concentration of chemical species R, W, R*, and W*. Partial differential equations (PDEs) governing the system can thus be expressed as:

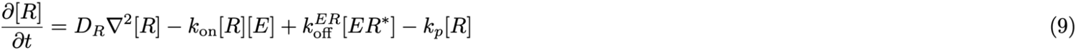

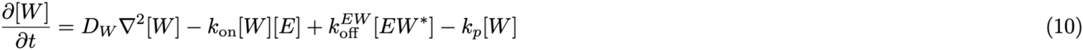

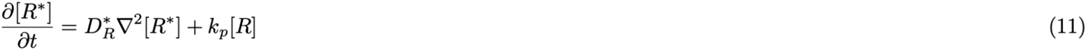

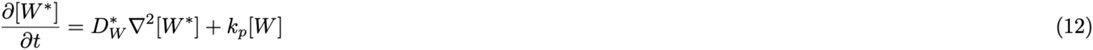

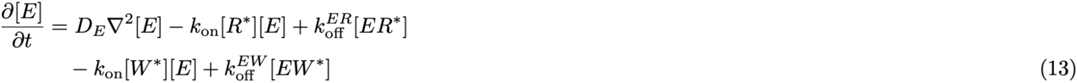

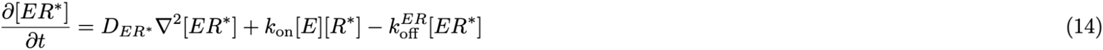

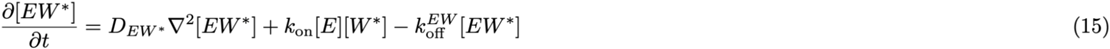

#### Selection of parameter values

Parameter values used for intracellular model analysis were selected from physiologically relevant values (Table S1). For example, the protein diffusion constant D is typically 0∼10 μm^2^/s, depending on different molecular masses in the cytoplasm.^28^ For simplicity, a uniform diffusion rate of 10 μm^2^/s was assumed across all species in the cytoplasm. Here, the association rate (k_on_) of the enzyme (E) and substrate (R* or W*) is set to 10^5^ M^-1^ s^-1^, which is within the typical k_on_ range for specific enzymes or enzyme classes.^67^ The dissociation rates of cognate (R*) and noncognate (W*) substrates are set to 0.1 s^-1^ and 1 s^-1^, respectively.^18^ The expected fidelity at equilibrium is hence η_eq_= k ^EW^ / k_off_ ^ER^ = 10. In our model, the kinases and phosphatases were assumed to act indiscriminately on both cognate and noncognate substrates. Both phosphorylation and dephosphorylation rate constants are in the range of 0.1∼100 s^-1^ as reported.^68,69^ The simulation used a cell diameter of 10 μm, approximating the typical diameter of human cells, such as HEK293 and HeLa cells.^28^

### Extracellular proofreading model coupled to endocytosis, self-activation, and mutual inhibition

#### Model framework

We developed a reaction-diffusion model to describe the dynamics of extracellular proofreading systems, including substrates R (cognate) and W (noncognate), the enzyme E, their complexes (ER and EW), and membrane-bound receptor-ligand complexes (R-receptor, W-receptor). The PDEs describing this system consist of diffusion terms and reaction terms. The diffusion term for each species C (where C here are [R], [W], [ER], [EW], and [E]) is expressed as D∇^2^C, with species-specific diffusion coefficients being D_R_, D_W_, D_ER_, D_EW_, and D_E_.

#### Boundary conditions and spatial configuration

The extracellular proofreading system forms quasi-one-dimensional (1D) concentration gradients of ER and EW. To model this, we define specific regions along the x-axis, with sender cells localized between 0 to 2000 μm, followed by proofreader cells occupying the adjacent region (x > 2000 μm). We imposed no-flux boundary conditions at both edges (x = 0 and x = L), ensuring the conservation of species, and continuity conditions in between. The discretized forms of these boundary conditions, using diffusion step size h, are:

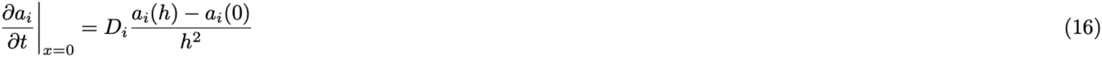

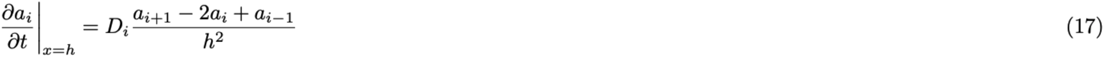

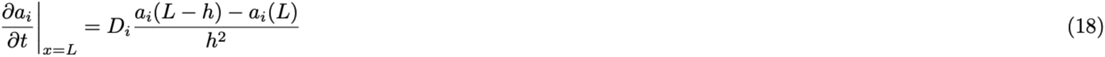

Here, a_i_ represents the concentration of species i (e.g., [ER], [EW]), and D_i_ is its corresponding diffusion coefficient.

#### Key biological processes

1. Secretion and receptor presentation. Sender cells constitutively secrete E, R, and W into the extracellular space at zero-order secretion rates J_Complex_. The synthesis rate for the receptor is J_Receptor_.
2. Enzyme-substrate binding. Substrates R and W bind to E with an association rate constant k_on_ and dissociation rate constants k_off_^ER^ and k_off_^EW^, respectively.
3. Receptor-substrate binding. R and W undergo reversible association and dissociation with membrane-bound receptors, with kinetics governed by their respective rate constants (k_on_, k_off_^receptor^).
4. Irreversible dissociation. Proofreader cells mediate endocytosis of receptor-substrate complexes (R-receptor and W-receptor) with a first-order rate constant γ.
5. Degradation. All species (R, W, E, ER, EW, R-receptor, W-receptor, receptor) are subject to first-order degradation with a constant rate of k_deg_, representing natural turnover in the extracellular environment.

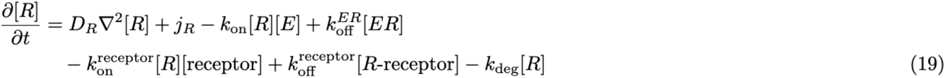

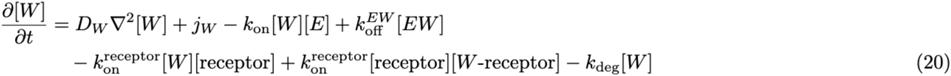

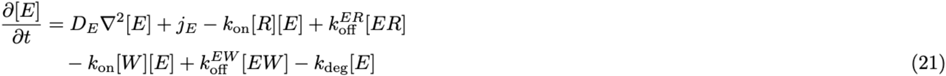

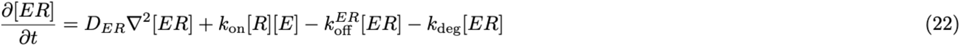

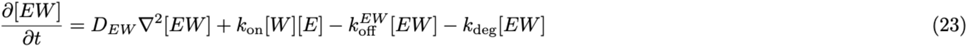

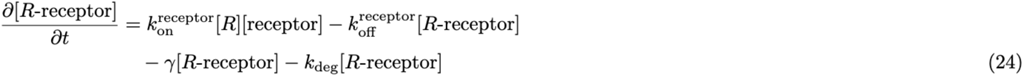

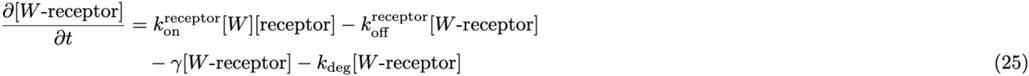

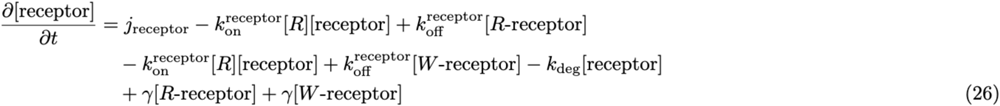

#### Modeling of self-activation, mutual inhibition, and SAMI

We assume self-activation, mutual inhibition, and SAMI follow the Hill function:

self-activation:

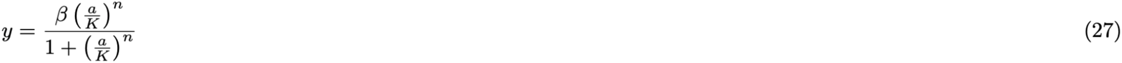

mutual inhibition:

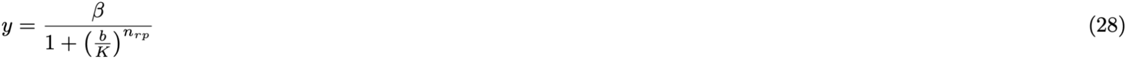

SAMI:

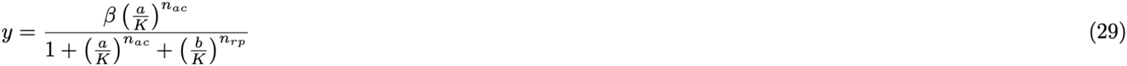

where,

- y is the protein secretion rate of proofreader cells
- β is the maximal secretion rate of the target protein by proofreader cells
- K here is the half-maximal effective concentration of the input protein
- n_ac_ is the Hill coefficient for self-activation
- n_rp_ is the Hill coefficient for mutual inhibition
- a, b are the concentrations of input proteins

The PDEs for ER and EW can be modified as:

(1) self-activation

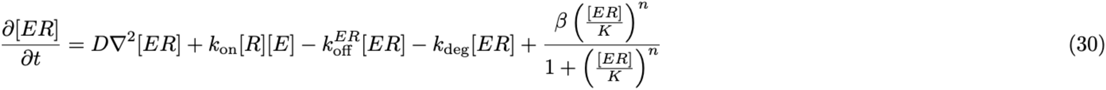

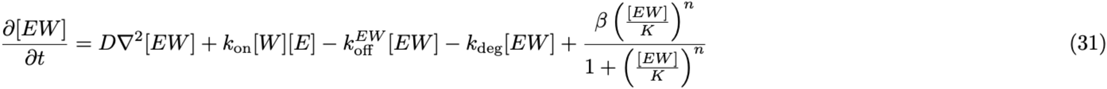

(2) mutual inhibition

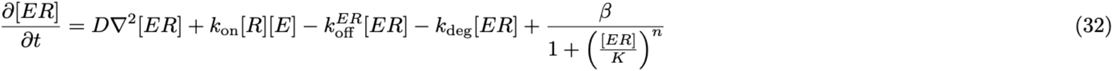

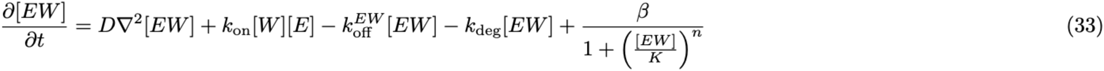

(3) SAMI

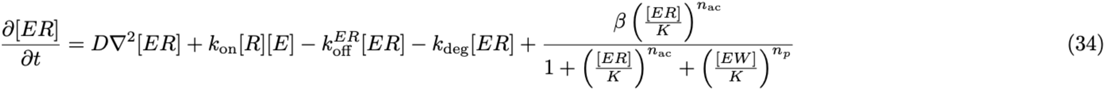

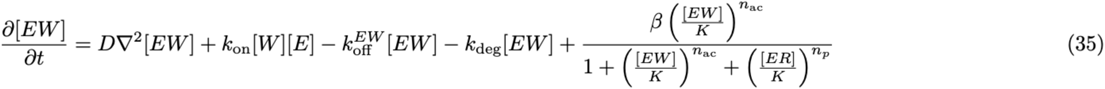

Selection of parameter values for the global parameter sweep

Parameter values used as the reference set for extracellular model analysis were selected based on a range of physiologically relevant values (Figure 2B). The protein diffusion rate in the extracellular space typically ranges from 0 to 50 μm^2^ s^-1^.^70–74^ The binding and dissociation rate constants (k_on_, k_off_^ER^, k_off_^EW^, and k_on_^receptor^) are consistent with the intercellular model, while the dissociation rate with endocytic receptors (k_off_^receptor^) ranges from 10^-5^ to 10^-2^ s^-1^.^75–77^ Here, the association rate (k_on_) of the enzyme (E) and substrate (R or W) is set to 10^-4^ nM^-1^ s^-1^ as the intracellular scheme.^67^ The endocytotic rate constant γ has been reported in the literature to range from 2.8 × 10^−4^ to 2.8 × 10^−3^ s^-1^,^29,78–82^ and we chose a wider range from 10^-5^ to 10^-2^ s^-1^. The secretion rate (J_Complex_) varies from 0 to 10^-2^ nM s^-1^,^29^ while the maximal rate for receptor synthesis (J_Receptor_) is similarly set at 10^-2^ s^-1^.^31^ The extracellular degradation rate is set at 2×10^−5^ s^-1^.^83,84^ These values serve as a reference set for the global parameter sweep to explore model behavior.

### Experimental design of the multi-molecular proofreading circuit

Principles established in the current work suggest the possibility of building a synthetic version of a proofreading circuit. This circuit could be constructed using a modular synthetic system centered on coiled-coil heterodimers as tunable substrates and enzymes, enabling precise control over cognate and noncognate binding affinities across a 10^-7^ to 10^-10^ M range, as confirmed by Surface Plasmon Resonance (SPR). To find model parameters such as diffusion constants, we utilize synthetic morphogen gradients (established via GFP-secreting sender and GFP-antibody-displaying receiver cells). The integration of clathrin-mediated endocytosis receptors will ensure rapid and irreversible internalization of dissociated ligands^85^. Synthetic receptors like SNIPR can sense ER and EW complexes,^86^ while post-translational regulations such as RELEASE could implement negative and positive feedback.^66,87^

#### Data and code availability

Raw data and simulation code are available at the GitHub repository https://github.com/BioProgramming-Lab/proofreading-2024-12 and Westalke datahub https://datahub.wllsb.edu.cn/#/datadetail/465a1f160d6a4ae6b08800bae493b954.

## ACKNOWLEDGMENTS

We thank Xiaoyu Wu for assistance in figure-making and theory development; Xinwen Fan, Xia Yao, Dingchen Yu, Wenhao Chen, and Xi Wang for helpful discussions on the project and suggestions on the manuscript; Westlake Center for Interdisciplinary Studies (CIS) for their helpful feedback on the project; the High-Performance Computing Center of Westlake University. This work was supported by the National Key R&D Program of China (2024YFA0919500), the Damon Runyon Cancer Research Foundation (DFS-63-24), and the Westlake Education Foundation.

## AUTHOR CONTRIBUTIONS

Conceptualization and investigation, Z.M., Y.J., F.X., and Z.C.; formal analysis, software, methodology, validation and visualization, Z.M., Y.J., Y.Y., B.W., F.X., and Z.C.; writing - original draft, Z.M., Y.J., and Z.C.; writing - review and editing, Z.M., Y.J., Y.Y., F.X., and Z.C.

**Table S1.** Parameters and their values used to model the intracellular spatial proofreading circuit.

| Parameter | Description | Value | Ref. |
| --- | --- | --- | --- |
| $k_{kin}$ | phosphorylation rate | $0.1 \sim 100 \text{s}^{-1}$ | 68,69 |
| $k_p$ | dephosphorylation rate | $0.1 \sim 100 \text{s}^{-1}$ | 68,69 |
| $k_{off}^{ER}$, $k_{off}^{EW}$ | dissociation rate constant | $0.1 \text{s}^{-1}$, $1 \text{s}^{-1}$ | 18 |
| D | diffusion rate | $0 - 10 \mu\text{m}^2 \text{s}^{-1}$ | 28 |
| Length | cell size | 10 $\mu\text{m}$ | 28 |

**Table S2.** Comparison of Single Molecular and Multi-Molecular KPR Systems.

| Category |  | Single Molecular | Multi-Molecular |
| --- | --- | --- | --- |
| Characteristics |  | fixed number of molecules, independent and identical molecules<sup>3,6,7</sup> | fluctuating number of molecules, interactions among molecules |
| Biological Example |  | enzymatic reactions<sup>9,14,51</sup>, T-cell activation<sup>8</sup> | spatial proofreading<sup>23</sup> |
| Performance Metrics | Accuracy | fidelity<sup>3,6,7</sup> | fidelity |
|  | Energy Consumption | number of intermediate states (n)<sup>16,48,88</sup> | endocytosis; auxiliary steps, e.g., SAMI |
|  | Speed | first hitting time of active state<sup>45</sup> | N/A |
|  | Activity | concentration of cognate complexes<sup>89</sup> | flux of cognate complexes |

**Table S3.** Parameters and their values used to model the classic kinetic proofreading circuit.

| Parameter | Description | Value |
| --- | --- | --- |
| $k_{\text{on}}$ | association rate of ER and EW | $10^{-6} \sim 10^6 \text{ nM s}^{-1}$ |
| $k_{\text{off}}^{\text{ER}}$ | dissociation rate of | $10^{-6} \sim 10^6 \text{ s}^{-1}$ |
| $k_{\text{off}}^{\text{EW}} / k_{\text{off}}^{\text{ER}}$ | ratio of dissociation rates between ER and EW | 1000 |
| $k_{\text{transit}}$ | transition rate | $10^{-6} \sim 10^6 \text{ s}^{-1}$ |
| $k_{\text{cat}}$ | the catalytic rate for product formation | $10^{-6} \sim 10^6 \text{ s}^{-1}$ |

\* The parameter values used here are out of the range of biologically plausible values to fully explore the boundary of activity-fidelity trade-off.

## REFERENCES

1. Lodish, H., Berk, A., Kaiser, C.A., Krieger, M., Bretscher, A., Ploegh, H., Amon, A., Martin, K.C. (2016). Molecular Cell Biology, 8th Edition (W. H. Freeman).

2. Loftfield, R.B., Hecht, L.I., and Eigner, E.A. (1963). The measurement of amino acid specificity of transfer ribonucleic acid. Biochim. Biophys. Acta: Nucleic Acids 72, 383–390.

3. Boeger, H. (2022). Kinetic proofreading. Annu. Rev. Biochem. 91, 423–447.

4. Alon, U. (2019). An introduction to systems biology: Design principles of biological circuits 2nd ed. (Chapman & Hall/CRC).

5. George, A.J.T., Stark, J., and Chan, C. (2005). Understanding specificity and sensitivity of T-cell recognition. Trends Immunol. 26, 653–659.

6. Hopfield, J.J. (1974). Kinetic proofreading: a new mechanism for reducing errors in biosynthetic processes requiring high specificity. Proc. Natl. Acad. Sci. U. S. A. 71, 4135–4139.

7. Ninio, J. (1975). Kinetic amplification of enzyme discrimination. Biochimie 57, 587–595.

8. McKeithan, T.W. (1995). Kinetic proofreading in T-cell receptor signal transduction. Proc. Natl. Acad. Sci. U. S. A. 92, 5042–5046.

9. Rodnina, M.V., and Wintermeyer, W. (2001). Fidelity of aminoacyl-tRNA selection on the ribosome: kinetic and structural mechanisms. Annu. Rev. Biochem. 70, 415–435.

10. Huang, W.Y.C., Alvarez, S., Kondo, Y., Lee, Y.K., Chung, J.K., Lam, H.Y.M., Biswas, K.H., Kuriyan, J., and Groves, J.T. (2019). A molecular assembly phase transition and kinetic proofreading modulate Ras activation by SOS. Science 363, 1098–1103.

11. Voisinne, G., Locard-Paulet, M., Froment, C., Maturin, E., Menoita, M.G., Girard, L., Mellado, V., Burlet-Schiltz, O., Malissen, B., Gonzalez de Peredo, A., et al. (2022). Kinetic proofreading through the multi-step activation of the ZAP70 kinase underlies early T cell ligand discrimination. Nat. Immunol. 23, 1355–1364.

12. Tischbirek, C.H., Colón, K.L., Lobbia, S., Cronin, C.J., and Cai, L. (2025). Spatial Proofreading Amplification of in situ Transcript and Protein Signals. Molecular Biology.

13. Morita, S., and Groves, J.T. (2025). Parallel reactions on a single T cell receptor offer a robust kinetic proofreading mechanism. Proc. Natl. Acad. Sci. U. S. A. 122, e2514057122.

14. Hopfield, J.J., Yamane, T., Yue, V., and Coutts, S.M. (1976). Direct experimental evidence for kinetic proofreading in amino acylation of tRNAIle. Proc. Natl. Acad. Sci. U. S. A. 73, 1164–1168.

15. Yamane, T., and Hopfield, J.J. (1977). Experimental evidence for kinetic proofreading in the aminoacylation of tRNA by synthetase. Proc. Natl. Acad. Sci. U. S. A. 74, 2246–2250.

16. Murugan, A., Huse, D.A., and Leibler, S. (2012). Speed, dissipation, and error in kinetic proofreading. Proc. Natl. Acad. Sci. U. S. A. 109, 12034–12039.

17. Wong, F., Amir, A., and Gunawardena, J. (2018). Energy-speed-accuracy relation in complex networks for biological discrimination. Phys. Rev. E 98, 012420.

18. Cui, W., and Mehta, P. (2018). Identifying feasible operating regimes for early T-cell recognition: The speed, energy, accuracy trade-off in kinetic proofreading and adaptive sorting. PLoS One 13, e0202331.

19. Hopfield, J.J. (1980). The energy relay: a proofreading scheme based on dynamic cooperativity and lacking all characteristic symptoms of kinetic proofreading in DNA replication and protein synthesis. Proc. Natl. Acad. Sci. U. S. A. 77, 5248–5252.

20. Buchel, G., Nayak, A.R., Herbine, K., Sarfallah, A., Sokolova, V.O., Zamudio-Ochoa, A., and Temiakov, D. (2023). Structural basis for DNA proofreading. Nat. Commun. 14, 8501.

21. Jenner, L., Demeshkina, N., Yusupova, G., and Yusupov, M. (2010). Structural rearrangements of the ribosome at the tRNA proofreading step. Nat. Struct. Mol. Biol. 17, 1072–1078.

22. Schamel, W.W., Risueño, R.M., Minguet, S., Ortíz, A.R., and Alarcón, B. (2006). A conformation-and avidity-based proofreading mechanism for the TCR–CD3 complex. Trends Immunol. 27, 176–182.

23. Galstyan, V., Husain, K., Xiao, F., Murugan, A., and Phillips, R. (2020). Proofreading through spatial gradients. eLife 9, e60415.

24. Kholodenko, B.N. (2006). Cell-signalling dynamics in time and space. Nat. Rev. Mol. Cell Biol. 7, 165–176.

25. Zabel, U., Schreck, R., and Baeuerle, P.A. (1991). DNA binding of purified transcription factor NF-kappa B. Affinity, specificity, Zn2+ dependence, and differential half-site recognition. J. Biol. Chem. 266, 252–260.

26. Garner, R.M., Molines, A.T., Theriot, J.A., and Chang, F. (2023). Vast heterogeneity in cytoplasmic diffusion rates revealed by nanorheology and Doppelgänger simulations. Biophys. J. 122, 767–783.

27. Chen, Y., Huang, J.-H., Phong, C., and Ferrell, J.E., Jr (2024). Viscosity-dependent control of protein synthesis and degradation. Nat. Commun. 15, 2149.

28. Milo, R., and Phillips, R. (2015). Cell biology by the numbers 1st Edition. (Garland Science) 10.1201/9780429258770.

29. Zhu, R., Santat, L.A., Markson, J.S., Nandagopal, N., Gregrowicz, J., and Elowitz, M.B. (2023). Reconstitution of morphogen shuttling circuits. Sci. Adv. 9, eadf9336.

30. Toda, S., McKeithan, W.L., Hakkinen, T.J., Lopez, P., Klein, O.D., and Lim, W.A. (2020). Engineering synthetic morphogen systems that can program multicellular patterning. Science 370, 327–331.

31. Stapornwongkul, K.S., de Gennes, M., Cocconi, L., Salbreux, G., and Vincent, J.-P. (2020). Patterning and growth control in vivo by an engineered GFP gradient. Science 370, 321–327.

32. Davis, S.J., Ikemizu, S., Evans, E.J., Fugger, L., Bakker, T.R., and van der Merwe, P.A. (2003). The nature of molecular recognition by T cells. Nat. Immunol. 4, 217–224.

33. Joedicke, L., Mao, J., Kuenze, G., Reinhart, C., Kalavacherla, T., Jonker, H.R.A., Richter, C., Schwalbe, H., Meiler, J., Preu, J., et al. (2018). The molecular basis of subtype selectivity of human kinin G-protein-coupled receptors. Nat. Chem. Biol. 14, 284–290.

34. Khodr, V., Machillot, P., Migliorini, E., Reiser, J.-B., and Picart, C. (June, 08 2021). High-throughput measurements of bone morphogenetic protein/bone morphogenetic protein receptor interactions using biolayer interferometry. Biointerphases 16, 031001.

35. Evans, C.R., Fan, Y., Weiss, K., and Ling, J. (2018). Errors during gene expression: Single-cell heterogeneity, stress resistance, and microbe-host interactions. MBio 9. 10.1128/mBio.01018-18.

36. Chen, Z., Linton, J.M., Xia, S., Fan, X., Yu, D., Wang, J., Zhu, R., and Elowitz, M.B. (2024). A synthetic protein-level neural network in mammalian cells. Science 386, 1243–1250.

37. Zhang, R., Goetz, H., Melendez-Alvarez, J., Li, J., Ding, T., Wang, X., and Tian, X.-J. (2021). Winner-takes-all resource competition redirects cascading cell fate transitions. Nat. Commun. 12, 1–9.

38. Chakravarty, S., Guttal, R., Zhang, R., and Tian, X.-J. (2024). Mitigating winner-take-all resource competition through antithetic control mechanism. ACS Synth. Biol. 13, 4050–4060.

39. Şimşek, E., Kim, K., Lu, J., Silver, A., Luo, N., Lee, C.T., and You, L. (2024). A “rich-get-richer” mechanism drives patchy dynamics and resistance evolution in antibiotic-treated bacteria. Mol. Syst. Biol. 20, 880–897.

40. Cherry, K.M., and Qian, L. (2018). Scaling up molecular pattern recognition with DNA-based winner-take-all neural networks. Nature 559, 370–376.

41. Yue, H.-Y., Bieberich, E., and Xu, J. (2017). Promotion of endocytosis efficiency through an ATP-independent mechanism at rat calyx of Held terminals. J. Physiol. 595, 5265–5284.

42. Kaksonen, M., and Roux, A. (2018). Mechanisms of clathrin-mediated endocytosis. Nat. Rev. Mol. Cell Biol. 19, 313–326.

43. Attwell, D., and Laughlin, S.B. (2001). An energy budget for signaling in the grey matter of the brain. J. Cereb. Blood Flow Metab. 21, 1133–1145.

44. Lynch, M., and Marinov, G.K. (2015). The bioenergetic costs of a gene. Proc. Natl. Acad. Sci. U. S. A. 112, 15690–15695.

45. Banerjee, K., Kolomeisky, A.B., and Igoshin, O.A. (2017). Elucidating interplay of speed and accuracy in biological error correction. Proc. Natl. Acad. Sci. U. S. A. 114, 5183–5188.

46. Bennett, C.H. (1979). Dissipation-error tradeoff in proofreading. Biosystems 11, 85–91.

47. Rao, R., and Peliti, L. (2015). Thermodynamics of accuracy in kinetic proofreading: dissipation and efficiency trade-offs. J. Stat. Mech: Theory Exp. 2015. 10.1088/1742-5468/2015/06/P06001.

48. Yu, Q., Kolomeisky, A.B., and Igoshin, O.A. (2022). The energy cost and optimal design of networks for biological discrimination. J. R. Soc. Interface 19, 20210883.

49. Piñeros, W.D., and Tlusty, T. (2020). Kinetic proofreading and the limits of thermodynamic uncertainty. Phys. Rev. E 101, 022415.

50. Chiuchiu, D., Mondal, S., and Pigolotti, S. (2023). Pareto optimal fronts of kinetic proofreading. New J. Phys. 25, 043007.

51. Kunkel, T.A., and Bebenek, K. (2000). DNA replication fidelity. Annu. Rev. Biochem. 69, 497–529.

52. Lever, M., Maini, P.K., van der Merwe, P.A., and Dushek, O. (2014). Phenotypic models of T cell activation. Nat. Rev. Immunol. 14, 619–629.

53. Johnson, K. (1993). Conformational Coupling in DNA Polymerase Fidelity. Annu. Rev. Biochem. 62, 685–713.

54. Gromadski, K.B., and Rodnina, M.V. (2004). Kinetic determinants of high-fidelity tRNA discrimination on the ribosome. Mol. Cell 13, 191–200.

