## supplemental figures+text S1+ text S22 for "Multimolecular proofreading overcomes the activity-fidelity trade-off"

A

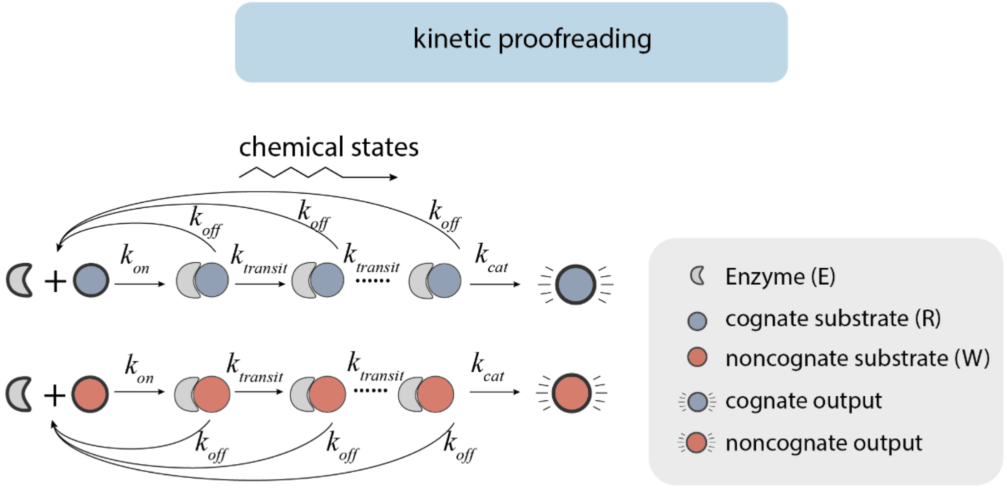


B

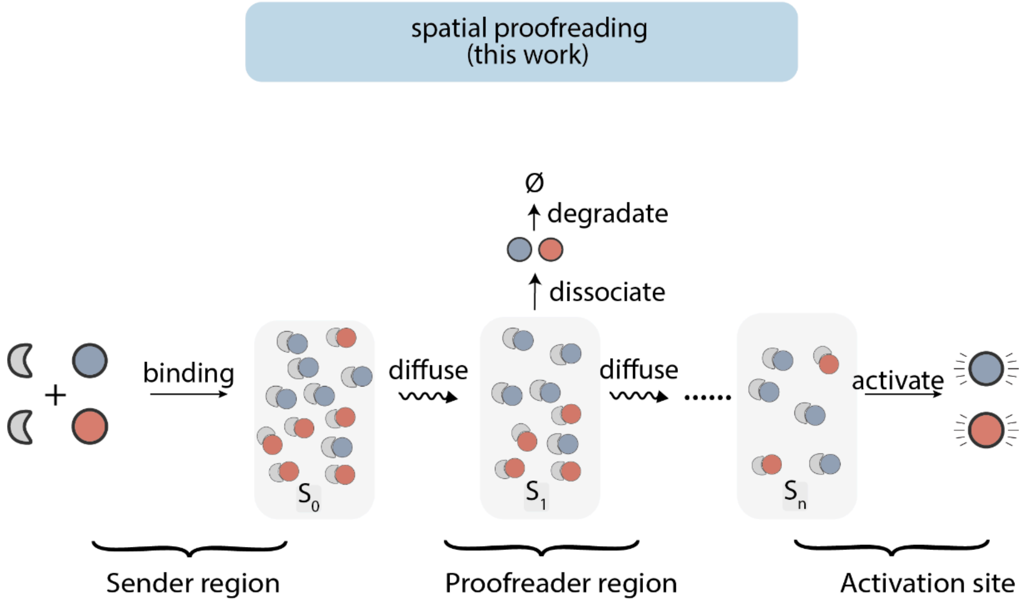

**Figure S1. Comparison between the single molecular T-cell activation KPR scheme and the multi-molecular, diffusion-based KPR**

(A) A representative kinetic proofreading model. Cognate and noncognate complexes diffuse from the sender region ($S_0$) to the proofreader region ($S_1$-$S_{n-1}$). During diffusion, these complexes could dissociate into monomers, and free substrates not bound by enzymes are endocytosed at the proofreader region, generating concentration gradients. At the activation site ($S_n$), located at the other end of the proofreader region, cognate and noncognate substrates are converted to their respective products.

(B) Schematics of the spatial proofreading system. Cognate and noncognate complexes bind and then diffuse from the sender region ($S_0$) to the proofreader region ($S_1$-$S_{n-1}$) with the diffusion rate of D. $k_{on,}$ and $k_{off}$ are the association constant and dissociation constants between enzymes (E) and cognate, or noncognate substrates (R or W). Free substrates not bound by enzymes are deactivated at a rate of γ at the proofreader region, generating concentration gradients. At the activation site ($S_n$), located at the other end of the proofreader region, cognate and noncognate substrates are converted to their respective products. $k_{cat}$ represents the catalytic rate for product formation.

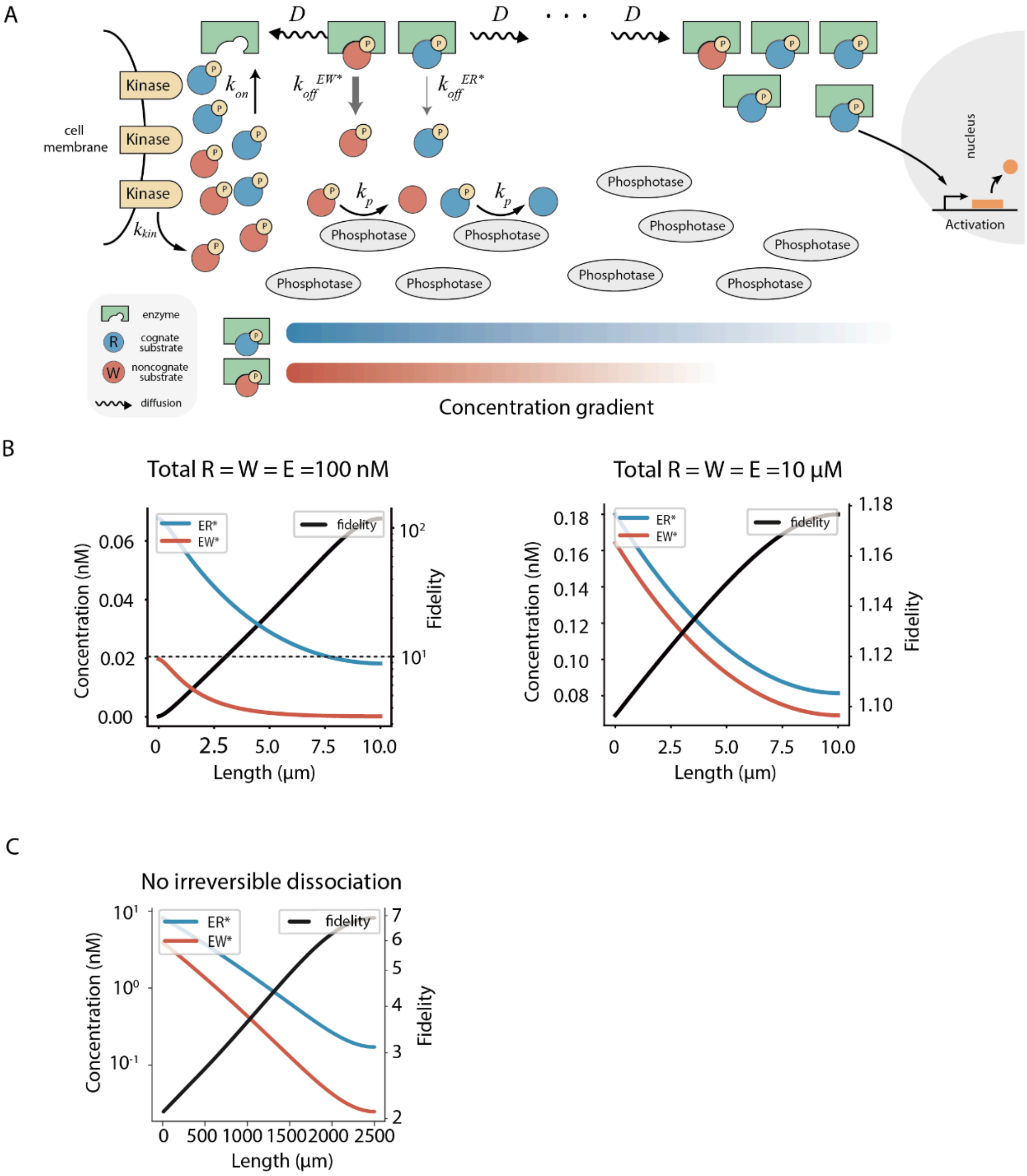

A
D
D
D
Kinase
Kinase
Kinase
cell membrane
$k_{on}$
$k_{off}^{EW*}$
$k_{off}^{ER*}$
$k_{kin}$
$k_p$
$k_p$
Phosphotase
Phosphotase
Phosphotase
Phosphotase
Phosphotase
Phosphotase
Phosphotase
Phosphotase
Phosphotase
Phosphotase
nucleus
Activation
enzyme
cognate substrate
noncognate substrate
diffusion
Concentration gradient
B
Total R = W = E =100 nM
ER*
EW*
fidelity
Concentration (nM)
Fidelity
Length (μm)
Total R = W = E =10 μM
ER*
EW*
fidelity
Concentration (nM)
Fidelity
Length (μm)
C
No irreversible dissociation
ER*
EW*
fidelity
Concentration (nM)
Fidelity
Length (μm)

**Figure S2. Analysis of intracellular and extracellular proofreading circuits**

(A) Schematic representation of the intracellular proofreading circuit. Cognate (blue) and noncognate (red) substrates are phosphorylated at the cell membrane by kinases at a rate of $k_{kin}$, and dephosphorylated in the cytosol at a rate of $k_p$. Phosphorylated substrates bind their enzymes with an association constant of $k_{on}$ and dissociation constants of $k_{off}^{ER}$ (cognate) and $k_{off}^{EW}$ (noncognate), respectively. Dissociated substrates are rapidly dephosphorylated and hence cannot rebind to the enzymes. Proteins diffuse in the cell with a diffusion constant of D. Complexes activate downstream signals upon reaching the nucleus. Parameter values used for following simulation are: $k_{kin}$ = 0.2 nM $s^{-1}$, $k_p$ = 5 $s^{-1}$, $k_{on}$ = 0.1 $nM^{-1}$ $s^{-1}$, $k_{off}^{ER}$ = 0.1 $s^{-1}$, $k_{off}^{EW}$ = 1 $s^{-1}$ and D = 1 $\mu m^2$ $s^{-1}$.
(B) Concentration gradients of ER* and EW* (blue and red, respectively; left axis), and the circuit fidelity (black; right axis) of a particular parameter combination. Fidelity is defined as the ratio of the concentrations of ER* to EW* at a given location. The difference in affinities between ER* and EW* complexes is 10-fold, and the circuit is considered to have achieved proofreading if the fidelity exceeds 10 (dashed line). The left and right panels represent simulations where the initial concentrations of E, R, and W are set to 100 nM and 10 μM, respectively.
(C) Concentration gradients of ER* and EW* (blue and red, respectively; left axis) and fidelity (black; right axis), in the absence of irreversible dissociation. Fidelity scales with distance but stays below 10, indicating no proofreading. Parameter values used for this simulation are: $J_{Complex}$ = $4\times10^{-4}$ nM $s^{-1}$, $J_{Receptor}$= 0 nM $s^{-1}$, $k_{off}^{receptor}$ = $4.5\times10^{-4}$ nM $s^{-1}$ $k_{off}^{ER}$ = $1\times10^{-4}$ $s^{-1}$, $k_{off}^{EW}$ = $4\times10^{-4}$ $s^{-1}$, and $\gamma$ = $4\times10^{-4}$ $s^{-1}$.

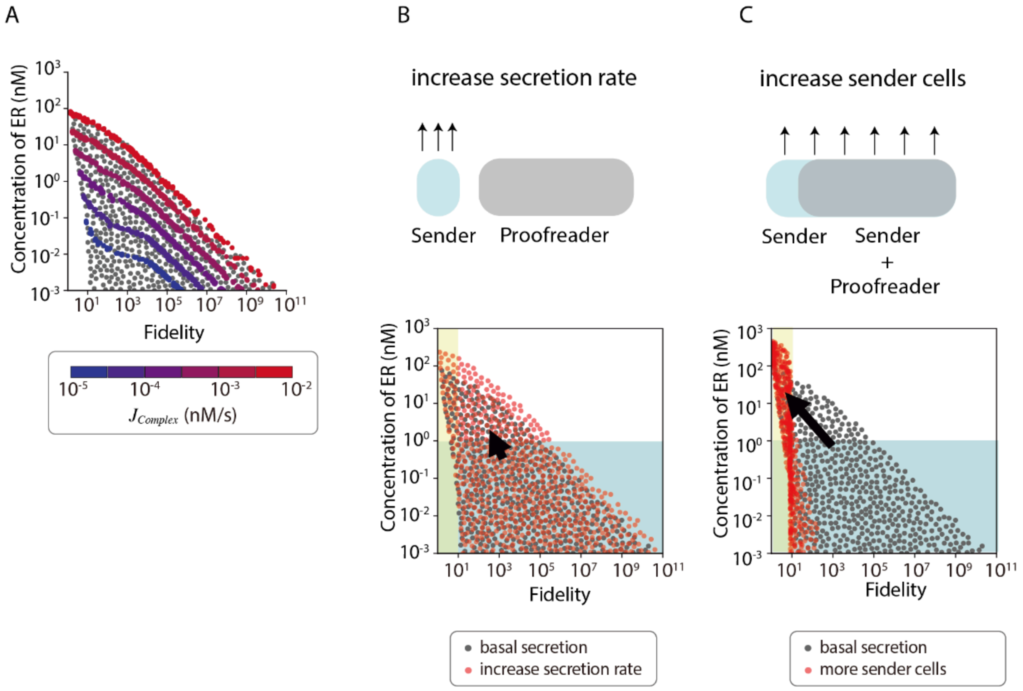
A
B
C
increase secretion rate
Sender
Proofreader
increase sender cells
Sender
Sender
+
Proofreader
Concentration of ER (nM)
Fidelity
$J_{Complex}$ (nM/s)
basal secretion
increase secretion rate
basal secretion
more sender cells

**Figure S3. Increasing secretion rates or the number of sender cells could only slightly affect the activity-fidelity trade-off**

(A) The secretion rates ($J_{Complex}$) primarily govern the boundary of activity-fidelity trade-off. The count-$J_{Complex}$ histogram shows that the number of parameter combinations, enabling proofreading, increases with higher $J_{Complex}$ values. The area-$J_{Complex}$ histogram in the bottom panel depicts the proofreading region spanned by parameter combinations in the activity-fidelity profile, which expands with elevated $J_{Complex}$ ranges. The proofreading region is where fidelity is higher than 10, and the final concentration of ER is less than 1 nM.

(B) Further increasing the secretion rates by sender cells does not significantly overcome the activity-fidelity trade-off. Red and gray dots represent increased and basal secretion rates by sender cells, respectively.

(C) Increasing the total number of sender cells does not overcome the activity-fidelity trade-off. An increase in concentration is accompanied by a significant decrease in fidelity. Black arrows indicate the overall migration trajectories of simulation outcomes.

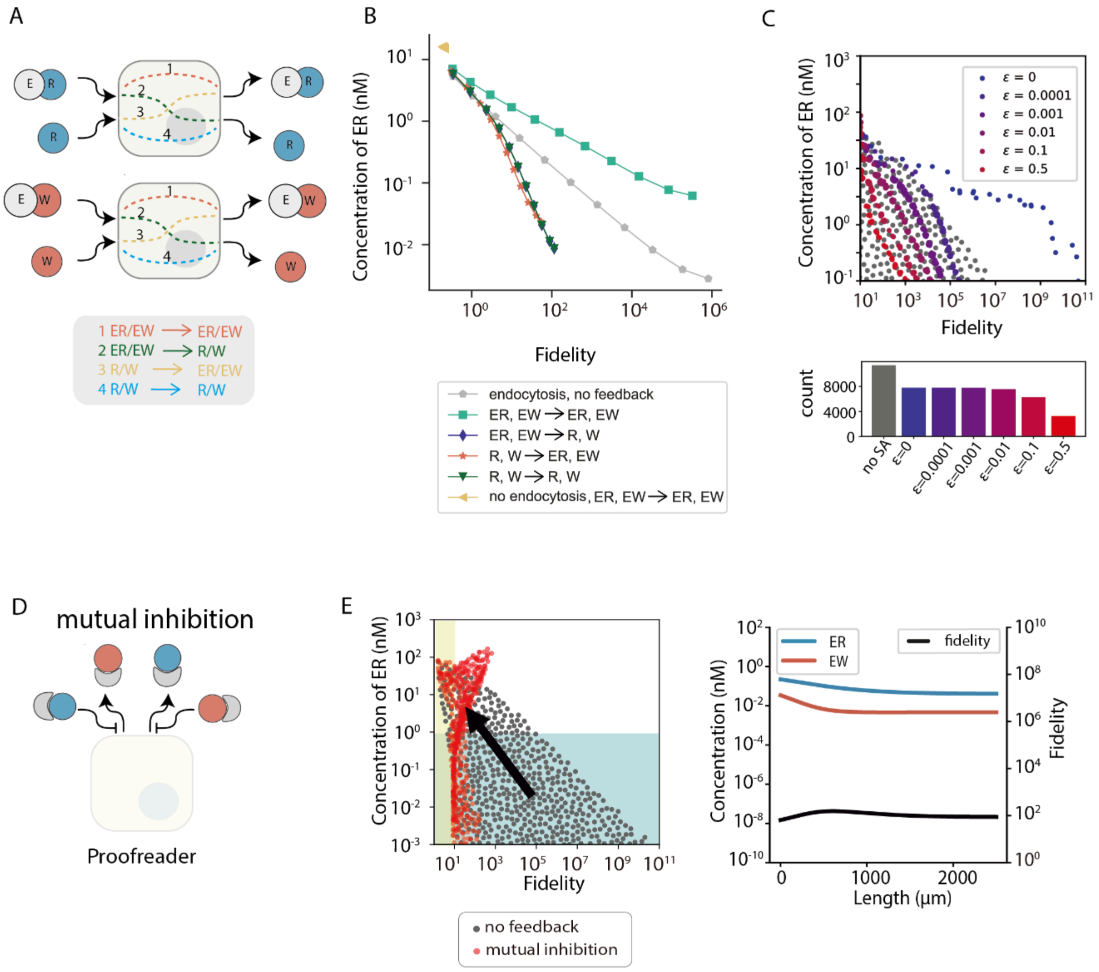
A
1 ER/EW → ER/EW
2 ER/EW → R/W
3 R/W → ER/EW
4 R/W → R/W
B
Concentration of ER (nM)
Fidelity
endocytosis, no feedback
ER, EW → ER, EW
ER, EW → R, W
R, W → ER, EW
R, W → R, W
no endocytosis, ER, EW → ER, EW
C
Concentration of ER (nM)
Fidelity
ε = 0
ε = 0.0001
ε = 0.001
ε = 0.01
ε = 0.1
ε = 0.5
count
no SA
ε=0
ε=0.0001
ε=0.001
ε=0.01
ε=0.1
ε=0.5
D
mutual inhibition
Proofreader
E
Concentration of ER (nM)
Fidelity
no feedback
mutual inhibition
ER
EW
fidelity
Concentration (nM)
Length (μm)
Fidelity

**Figure S4. Various other schemes that could break the fidelity-activity trade-off**

(A) Schematics of self-activation mechanisms in proofreading circuits, showing four possible pairings of activators and secreted proteins: (1) ER / EW activating ER / EW, (2) ER / EW activating R / W, (3) R / W activating ER / EW, and (4) R / W activating R / W. Parameter values used for this simulation are: $J_{Complex}$ = 4×10$^{-4}$ nM s$^{-1}$, $J_{Receptor}$ = 2×10$^{-4}$ nM s$^{-1}$, $k_{off}^{receptor}$ = 4.5×10$^{-4}$ nM s$^{-1}$, $k_{off}^{ER}$ = 1×10$^{-4}$ s$^{-1}$, $k_{off}^{EW}$ = 4×10$^{-4}$ s$^{-1}$, γ = 4×10$^{-4}$ s$^{-1}$, and $k_{deg}$ = 2×10$^{-5}$ nM s$^{-1}$. . For those with feedback, the n = 1, K = 1, β = 1.

(B) Simulation results for the concentrations of ER under the four self-activation schemes, compared with no self-activation.

(C) Activity-fidelity trade-off for the SA circuit across various error rates. The scatter plot shows the activity-fidelity trade-off boundary for a range of error rates (ε) in the SA circuit: ε = 0, ε = 0.0001, ε = 0.001, ε = 0.01, ε = 0.1. Each color dot line represents the trade-off boundary for the corresponding error rate. The bar plot shows the count of simulation trajectories capable of proofreading in each error rate category.

(D) Schematic of mutual inhibition in proofreader cells. Proofreader cells endocytose dissociated substrates (R and W, respectively) for degradation and can secrete ER and EW at a basal secretion rate. At the same time, ER and EW can inhibit each other's synthesis and secretion.

(E) Simulation results of the extracellular proofreading circuit with (red) and without (gray) mutual inhibition. The yellow region, where fidelity is less than 10, indicates no proofreading. The blue region, where the final concentration of ER is less than 1 nM, indicates that the ER concentration is too low to elicit downstream signaling. The right panel represents concentration gradients of ER and EW (blue and red, respectively; left axis), and the circuit fidelity (black; right axis) of a parameter combination. The black arrow indicates the overall migration trajectory of simulation outcomes after incorporating mutual inhibition.

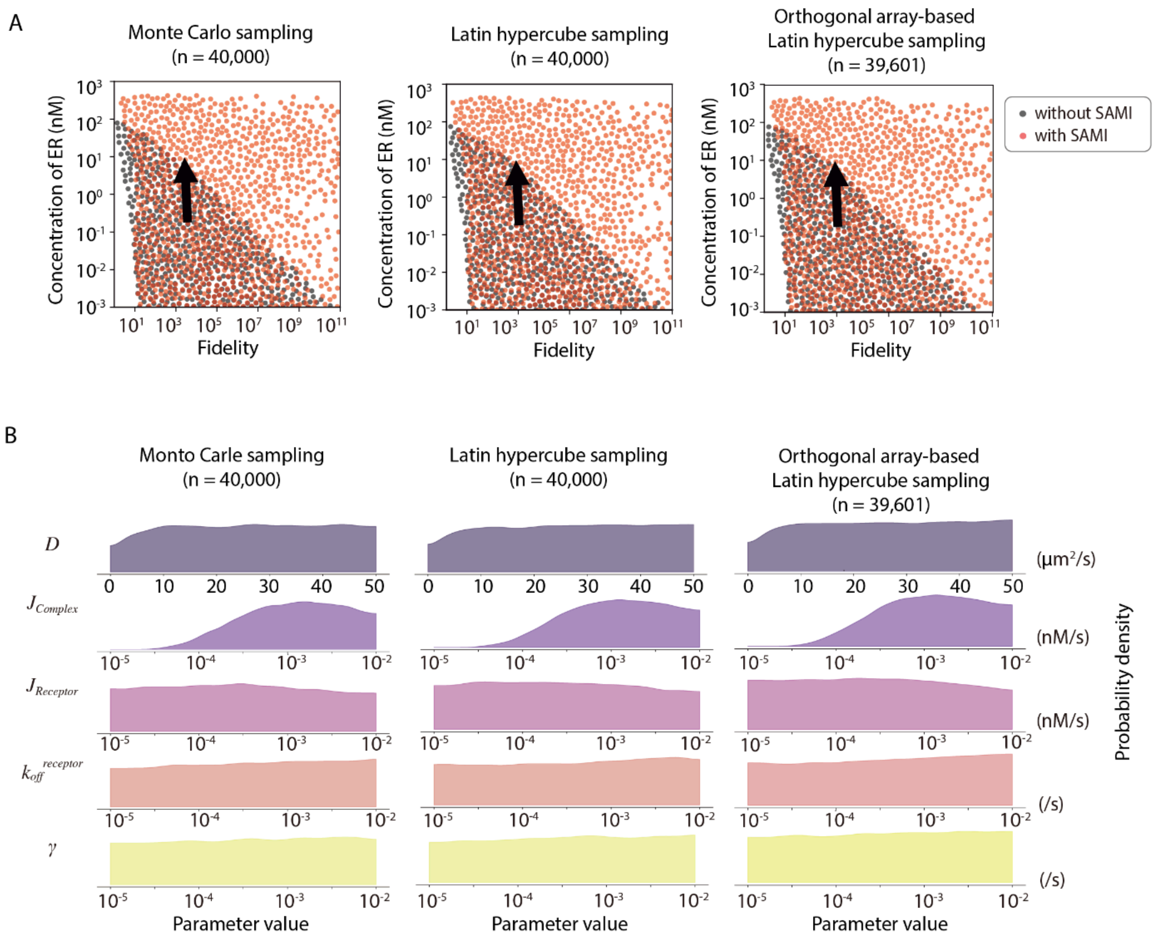
A
Monte Carlo sampling
(n = 40,000)
Latin hypercube sampling
(n = 40,000)
Orthogonal array-based
Latin hypercube sampling
(n = 39,601)
Concentration of ER (nM)
Fidelity
without SAMI
with SAMI
B
Monto Carle sampling
(n = 40,000)
Latin hypercube sampling
(n = 40,000)
Orthogonal array-based
Latin hypercube sampling
(n = 39,601)
D
J Complex
J Receptor
k off receptor
γ
(µm²/s)
(nM/s)
(nM/s)
(/s)
(/s)
Probability density
Parameter value

**Figure S5. Comparison among different sampling methods for the extracellular proofreading circuit.**

(A) Each dot represents the fidelity and ER concentration at the activation site for a parameter combination across the range of parameter values in Figure 2B.
(B) Designability plot showing parameter distributions that give rise to simulation trajectories within the red triangle in Figure 2C: from left to right are Monte Carlo sampling (MCS), Latin hypercube sampling (LHS), and orthogonal array-based Latin hypercube sampling. Parameter values used for this simulation are shown in Figure 2B.

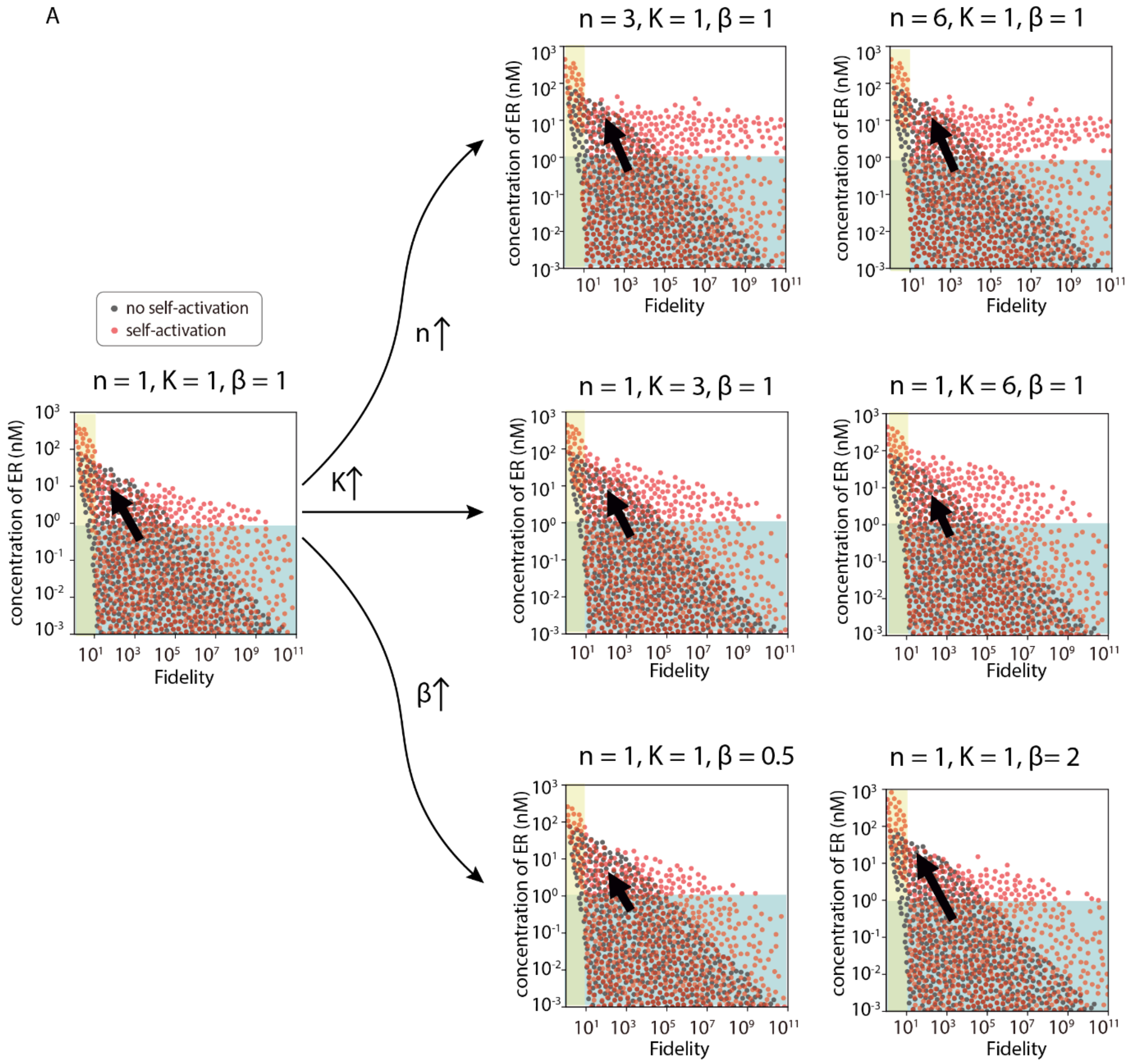

A
no self-activation
self-activation
n = 1, K = 1, β = 1
n↑
K↑
β↑
n = 3, K = 1, β = 1
n = 6, K = 1, β = 1
n = 1, K = 3, β = 1
n = 1, K = 6, β = 1
n = 1, K = 1, β = 0.5
n = 1, K = 1, β= 2
concentration of ER (nM)
Fidelity

**Figure S6. Various degrees of self-activation can tune the activity-fidelity trade-off**

(A) Simulation results of the extracellular proofreading circuit without (gray dots) and with self-activation (red dots) of the baseline parameter set ($n = 1$, $K = 1$, $\beta = 1$) for the activation Hill function. The effects of varying self-activation parameters on the activity-fidelity trade-off are plotted on the right. Each panel illustrates how changes in $\beta$ (the maximal secretion rate of the target protein by proofreader cells), $K$ (the half-maximal effective concentration of the input protein for self-activation), and $n$ (Hill coefficient) can affect the trade-off. Black arrows indicate overall migration trajectories of simulation outcomes.

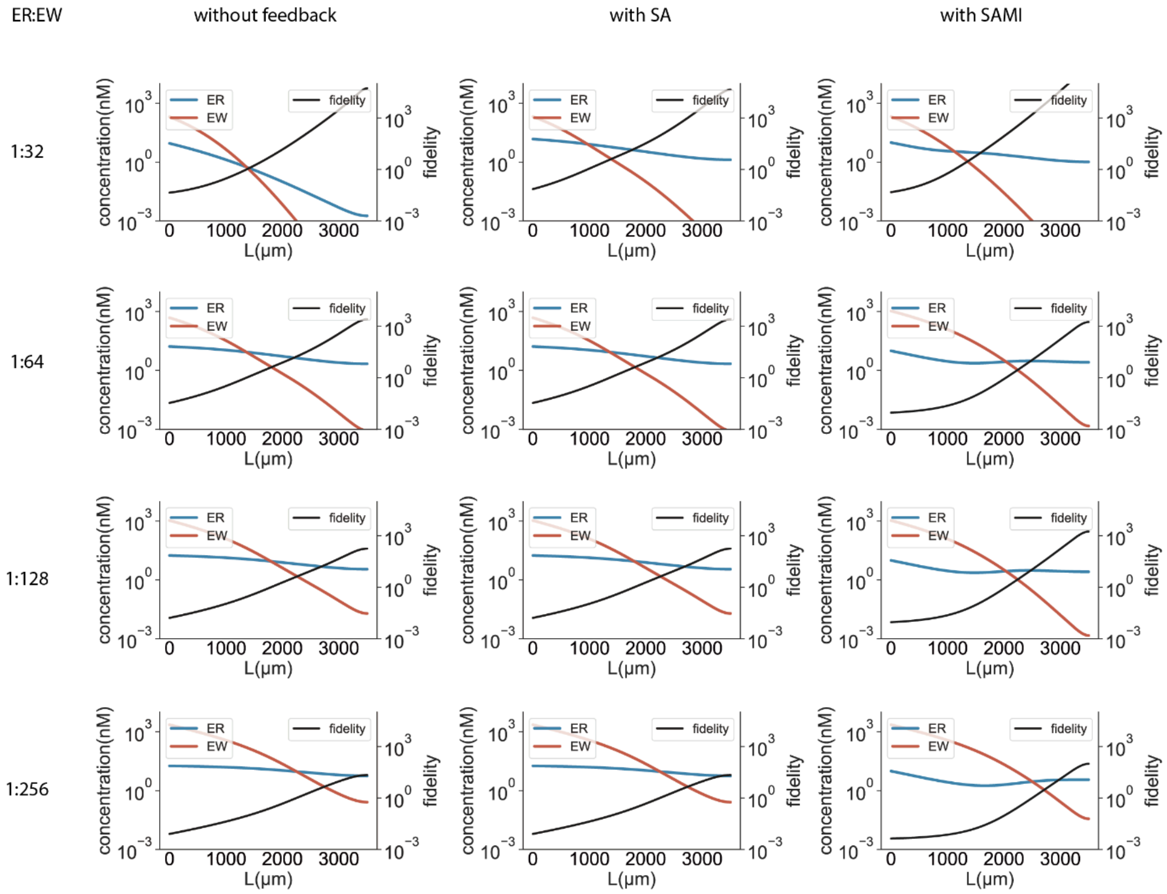

ER:EW
without feedback
with SA
with SAMI
1:32
1:64
1:128
1:256
ER
EW
fidelity
concentration(nM)
fidelity
L(μm)
0
1000
2000
3000
$10^3$
$10^0$
$10^{-3}$

**Figure S7. Effect of unequal ER and EW expression on proofreading without feedback, with SA, and with SAMI**

Concentration gradients of ER and EW (blue and red, respectively; left axis) and circuit fidelity (black; right axis) for a particular parameter combination are shown, both with and without feedback. Fidelity is defined as the ratio of the concentrations of ER to EW at a given location. Parameter values used for this simulation are: $J_{Complex}^{EW} = 2^{i} * J_{Complex}^{ER}$ (i = 4, 5, 6, or, 7), $J_{Complex}^{ER} = 4\times10^{-4}$ nM $s^{-1}$, $J_{Receptor} = 2\times10^{-4}$ nM $s^{-1}$, $k_{on}^{receptor} = 4.5\times10^{-4}$ nM $s^{-1}$, $k_{off}^{receptor} = 1\times10^{-3}$ $s^{-1}$, $k_{on}^{ER} = k_{on}^{EW} = 1\times10^{-4}$ nM $s^{-1}$, $k_{off}^{ER} = 1\times10^{-4}$ $s^{-1}$, $k_{off}^{EW} = 1\times10^{-3}$ $s^{-1}$, $\gamma = 4\times10^{-4}$ $s^{-1}$, and $k_{deg} = 2\times10^{-5}$ nM $s^{-1}$. For the Hill function of self-activation and mutual inhibition: k = 1 and n = 1.

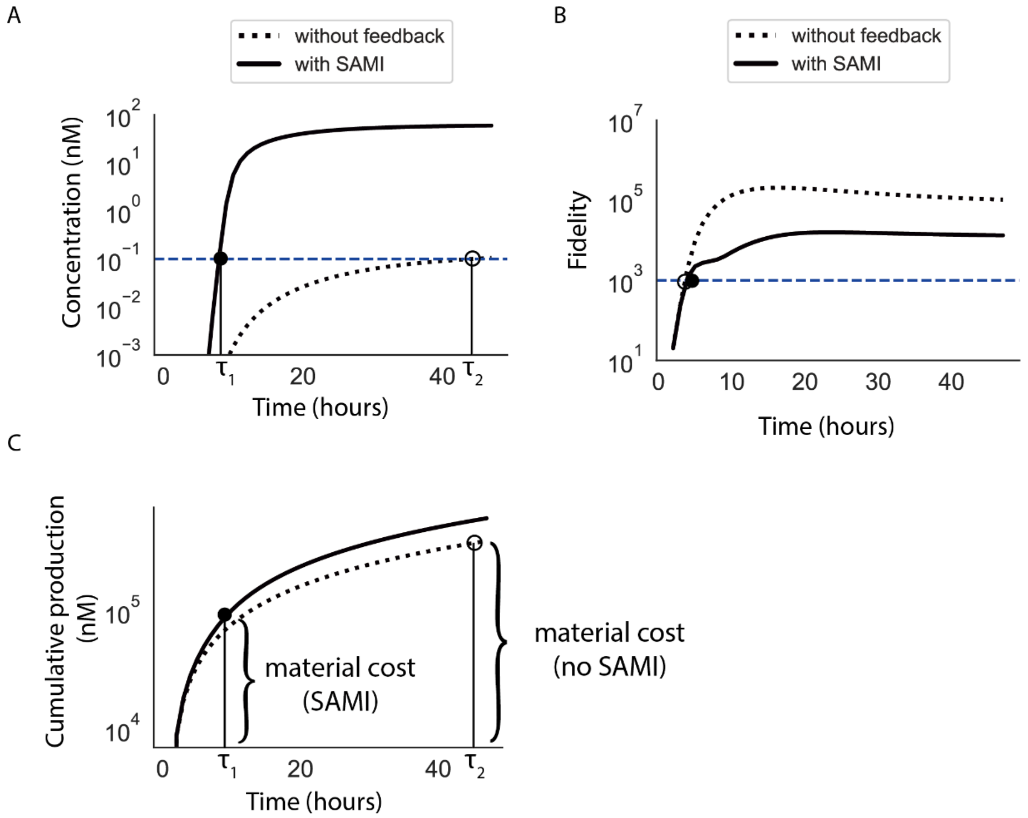

A
without feedback
with SAMI
Concentration (nM)
Time (hours)
B
without feedback
with SAMI
Fidelity
Time (hours)
C
Cumulative production (nM)
material cost (SAMI)
material cost (no SAMI)
Time (hours)

**Figure S8. Time and material cost analysis of spatial proofreading with and without SAMI**

(A) Dynamics of the ER concentration at a distance L = 2000 μm. Parameter values used for this simulation (also for B and C) are: $J_{Complex} = 3\times10^{-3}$ nM $s^{-1}$ (equal to the maximum secretion rate in positive/negative feedback), $J_{Receptor} = 2\times10^{-4}$ nM $s^{-1}$, $k_{on}^{receptor} = 4.5\times10^{-4}$ nM $s^{-1}$, $k_{off}^{receptor} = 1\times10^{-3}$ $s^{-1}$, $k_{on}^{ER} = k_{on}^{EW} = 1\times10^{-4}$ nM $s^{-1}$, $k_{off}^{ER} = 1\times10^{-4}$ $s^{-1}$, $k_{off}^{EW} = 1\times10^{-3}$ $s^{-1}$, $\gamma = 4\times10^{-4}$ $s^{-1}$, and $k_{deg} = 2\times10^{-5}$ nM $s^{-1}$, for the Hill function of self-activation and mutual inhibition: k = 1 and n = 1. Solid and dotted lines represent the concentration of ER complexes at the activation site (L = 2000 μm) with and without SAMI, respectively. The threshold shown by the blue dotted line corresponds to a concentration of 0.1 nM. The solid and hollow dots represent time points at which the concentration of ER complexes at the activation site reaches the activity threshold, with and without SAMI, respectively.

(B) Dynamics of the fidelity at a distance L = 2000 μm. Solid and dotted lines represent the fidelity at the activation site (L = 2000 μm) with and without SAMI, respectively. The threshold shown by the blue dotted line is a fidelity of $10^3$. The solid and hollow dots represent time points when the fidelity at the activation site reaches the fidelity threshold with and without SAMI, respectively.

(C) Cumulative production of E, R, and W molecules over time, with (solid line) and without (dotted line) SAMI.

A

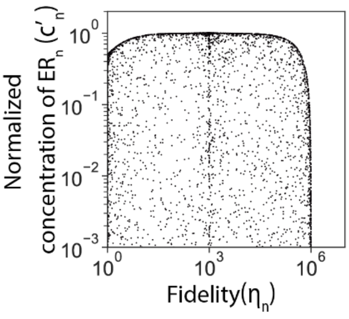

Normalized
concentration of $ER_n$ ($c'_n$)
$10^0$
$10^{-1}$
$10^{-2}$
$10^{-3}$
$10^0$
$10^3$
$10^6$
Fidelity($\eta_n$)

**Figure S9. The activity-fidelity trade-off persists with varying numbers of proofreading steps**

(A) Activity-fidelity trade-off of the KPR circuit with one intermediate state (n = 1) where R and W compete for enzyme binding. The normalized concentration of ER is $c'_n$, normalized by the total concentration of enzymes $c_0$ (free enzymes plus enzyme-substrate complexes), i.e. $c'_n = c_n / c_0$. Black dots represent simulation outcomes with various parameter value combinations listed in Table S3.

A

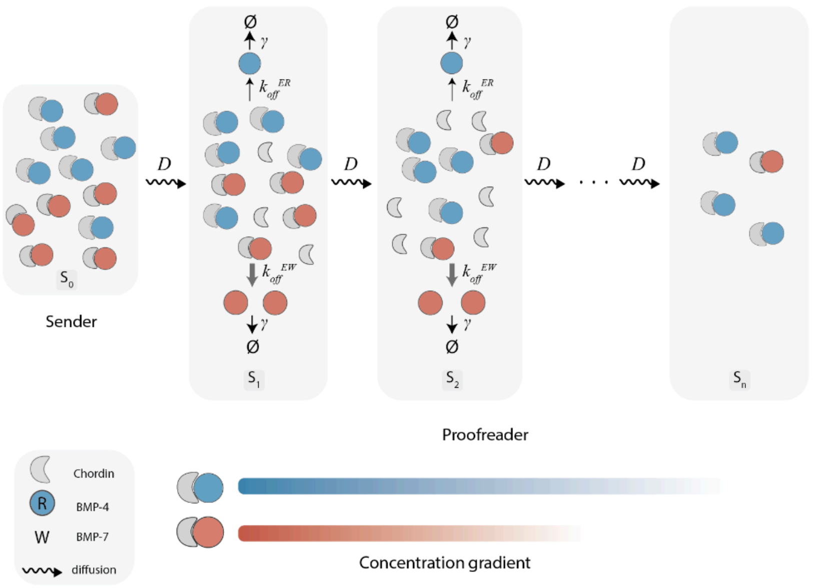


B

| parameter | description | value | ref. |
|---|---|---|---|
| $\gamma$ | endocytosis rate | $4\times10^{-4}$ s$^{-1}$ | [30,79-83] |
| $J_{Complex}$ | production rate of ER and EW | $4\times10^{-4}$ nM s$^{-1}$ | [30] |
| $J_{Receptor}$ | production rate of endocytic receptors | $2\times10^{-4}$ nM s$^{-1}$ | [32] |
| $k_{off}^{receptor}/k_{on}^{receptor}$ | dissociation and assocition rates between endocytic receptor and its target | $4.5\times10^{-4}$ nM s$^{-1}$<br>$1\times10^{-3}$ s$^{-1}$ | [76-78] |
| $D$ | diffusion rate | 10 μm$^2$ s$^{-1}$ | [71-75] |
| $k_{off}/k_{on}^{chd/BMP\text{-}4}$ | dissociation and assocition rates between Chordin and BMP-4 | $0.5\times10^{-4}$ s$^{-1}$<br>$3.9\times10^{-5}$ nM s$^{-1}$ | [58-60] |
| $k_{off}/k_{on}^{chd/BMP\text{-}7}$ | dissociation and assocition rates between Chordin and BMP-7 | $5.1\times10^{-4}$ s$^{-1}$<br>$2.3\times10^{-5}$ nM s$^{-1}$ | [58-60] |

C

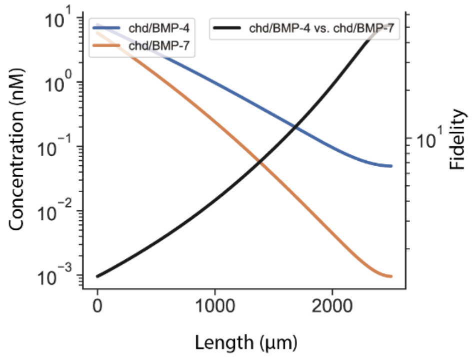

**Figure S10. Possible Spatial Proofreading in Morphogen Systems**

(A) Schematic of a potential spatial proofreading circuit in the BMP signaling network. Chordin/BMP-4 and Chordin/BMP-7 are considered to be the cognate and noncognate complexes (ER and EW). Cognate and noncognate complexes diffuse from the sender region ($S_0$) to the proofreader region ($S_1$-$S_{n-1}$). Free substrates not bound by enzymes are removed at the proofreader region, generating concentration gradients. At the activation site ($S_n$), located at the other end of the proofreader region, cognate and noncognate substrates are converted to their respective products.
(B) The parameters used for this simulation.
(C) Concentration gradients of Chrodin/BMP-4 and Chordin/BMP-7 (blue and red, respectively; left axis), and the circuit fidelity of a particular parameter combination (black; right axis). Fidelity is defined as the ratio of the concentrations of ER to EW at a given location. The difference in affinities between ER and EW complexes is slight, and the circuit is considered to have achieved proofreading as the fidelity constantly increases during diffusion.

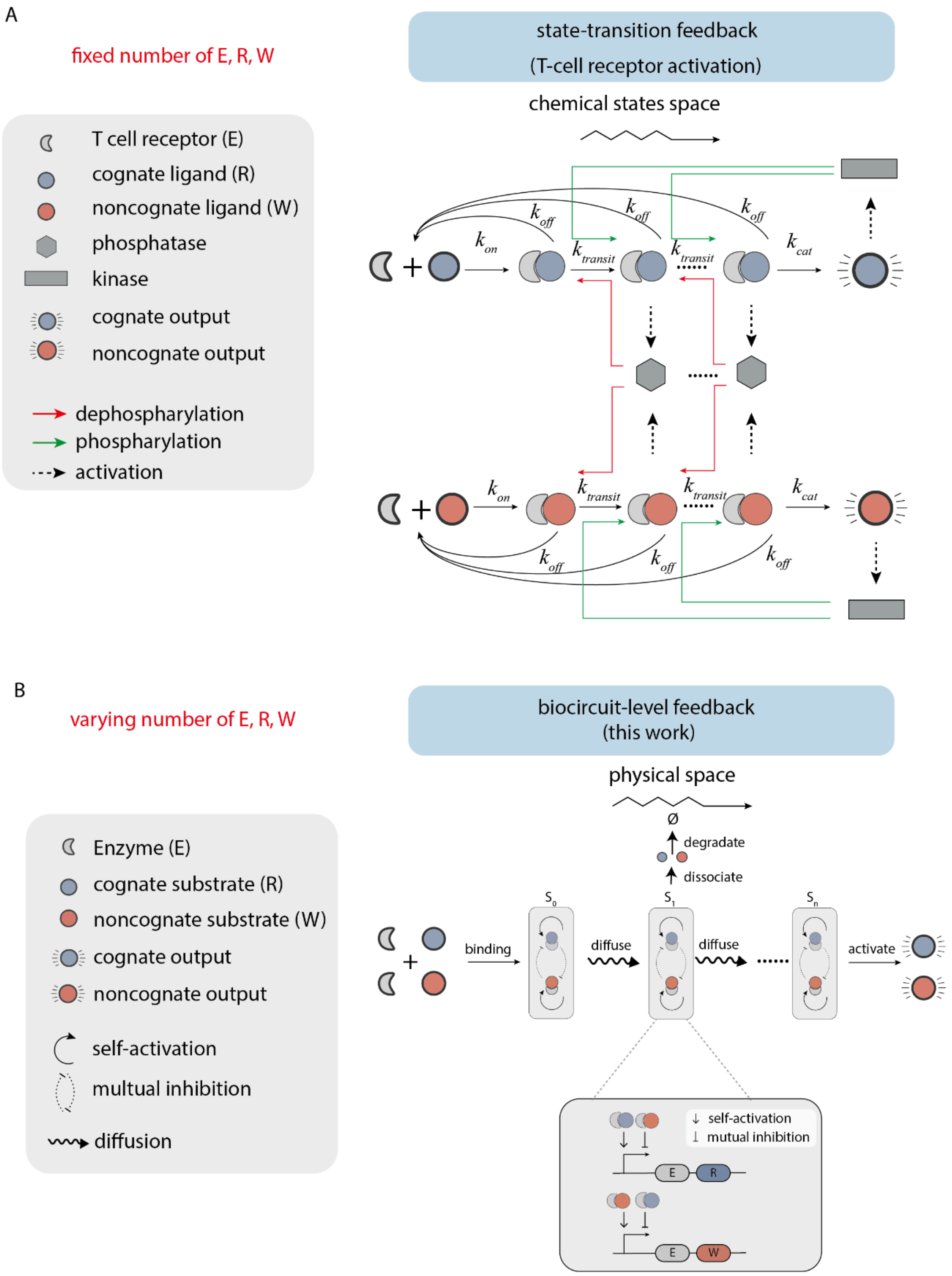
A
fixed number of E, R, W
state-transition feedback
(T-cell receptor activation)
chemical states space
T cell receptor (E)
cognate ligand (R)
noncognate ligand (W)
phosphatase
kinase
cognate output
noncognate output
dephosphorylation
phosphorylation
activation
$k_{on}$
$k_{off}$
$k_{transit}$
$k_{cat}$
B
varying number of E, R, W
biocircuit-level feedback
(this work)
physical space
Ø
degradate
dissociate
Enzyme (E)
cognate substrate (R)
noncognate substrate (W)
cognate output
noncognate output
self-activation
multual inhibition
diffusion
binding
diffuse
diffuse
activate
$S_0$
$S_1$
$S_n$
self-activation
mutual inhibition
E
R
E
W

**Figure S11. Comparison between the single molecular T-cell activation KPR scheme and the multi-molecular, diffusion-based KPR**

(A) A representative T-cell activation kinetic-proofreading model with positive and negative feedback loops. Cognate and noncognate ligands bind to T cell receptors (TCRs), leading to their phosphorylation. Phosphorylated TCR complexes activate phosphatases, which dephosphorylate the TCR complexes, thereby generating a negative feedback loop. Fully phosphorylated TCR activates kinases, which could add phospho groups to intermediate TCR complexes, thereby generating a positive feedback. The total number of molecules remains the same, as only single molecular state transitions are involved.
(B) Schematic of the spatial proofreading system with self-activation and mutual inhibition. Cognate and noncognate complexes diffuse from the sender region ($S_0$) to the proofreader region ($S_1$-$S_{n-1}$). During diffusion, these complexes could dissociate into monomers, and free substrates not bound by enzymes are endocytosed at the proofreader region, generating concentration gradients. At the activation site ($S_n$), located at the other end of the proofreader region, cognate and noncognate substrates are converted to their respective products. Proofreader cells possess additional receptors that sense the presence of ER and EW complexes and respond by synthesizing and secreting ER and EW molecules, respectively. Meanwhile, ER and EW can mutually inhibit their synthesis and secretion.

### Theoretical analysis of the activity-fidelity trade-off in classic kinetic proofreading systems

#### Description of the kinetic proofreading system

As described in Figure 5A of the main text, the classic kinetic proofreading system involves enzymes (E), cognate substrates (R), noncognate substrates (W), and enzyme-substrate complexes (EW and ER). Here, we provide the derivation of the performance of proofreading as well as the theoretical boundary of the activity-fidelity trade-off, as shown in Figure 5C.

We study the dynamics of kinetic proofreading reactions where the enzyme (E) binds to both cognate (R) and noncognate (W) substrates with the same association rate constant, $k_{\text{on}}$ and dissociation rate constants $k_{\text{off}}^{\text{ER}}$ or $k_{\text{off}}^{\text{EW}}$, respectively, forming the ER and EW complexes.

Here, $n$ denotes the number of kinetic proofreading steps, and the ER and EW complexes each have $n$ states, labeled ER$_i$ or EW$_i$ for $i = 0, 1, ..., n$. $k_{\text{transit}}$ denotes the transition rate for the kinetic proofreading steps. The final intermediate states ER$_n$ and EW$_n$ produce final products with the catalytic rate of $k_{\text{cat}}$. The chemical reactions governing the system are as follows:

$$\text{E} + \text{R} \underset{k_{\text{off}}^{\text{ER}}}{\overset{k_{\text{on}}}{\rightleftharpoons}} \text{ER}_0,$$

$$\text{E} + \text{W} \underset{k_{\text{off}}^{\text{EW}}}{\overset{k_{\text{on}}}{\rightleftharpoons}} \text{EW}_0,$$

$$\text{ER}_{\text{i}-1} \xrightarrow{k_{\text{transit}}} \text{ER}_{\text{i}},$$

$$\text{EW}_{\text{i}-1} \xrightarrow{k_{\text{transit}}} \text{EW}_{\text{i}},$$

$$\text{ER}_{\text{i}} \xrightarrow{k_{\text{off}}} \text{E} + \text{R},$$

$$\text{EW}_{\text{i}} \xrightarrow{k_{\text{off}}} \text{E} + \text{W},$$

$$\text{ER}_{\text{n}} \xrightarrow{k_{\text{cat}}} \text{E} + \text{P}_{\text{R}},$$

$$\text{EW}_{\text{n}} \xrightarrow{k_{\text{cat}}} \text{E} + \text{P}_{\text{W}}.$$

We simplify the system by neglecting the competition between E and W to independently study the proofreading effect. We denote a generic substrate by S, where $S \in \{R, W\}$. The concentrations of free substrate, enzyme, and the enzyme-substrate complexes are represented by $x_S$, $x_E$, $x_{ES,i}$ (with i = 0, 1, 2, ..., n), respectively.

The rate equations for the chemical reactions are given by:

$$\begin{aligned}
\frac{d}{dt}x_E &= -k_{\text{on}}x_E x_S + k_{\text{off}}^{\text{ES}} x_{ES,i} + (k_{\text{off}}^{\text{ES}} + k_{\text{cat}})x_{ES,n} \\
\frac{d}{dt}x_{ES,0} &= k_{\text{on}}x_E x_S - (k_{\text{off}}^{\text{ES}} + k_{\text{transit}})x_{ES,0} \\
\frac{d}{dt}x_{ES,i} &= k_{\text{transit}}x_{ES,i-1} - (k_{\text{off}}^{\text{ES}} + k_{\text{transit}})x_{ES,i}(i = 0, 1, 2, ..., n-1) \\
\frac{d}{dt}x_{ES,n} &= k_{\text{transit}}x_{ES,n-1} - (k_{\text{off}}^{\text{ES}} + k_{\text{cat}}^{\text{ES}})x_{ES,n}
\end{aligned}$$

Here, a simplifying assumption is made for overabundant substrate $S$, which leads to a linear system in terms of $(x_E, x_{ES,i})$. Under this assumption, the rate equations can be expressed as:

$$\frac{d}{dt}x = Mx.$$

To investigate the performance of kinetic proofreading, we focus on the steady-state behavior of the kinetic proofreading system. At steady state, the following condition holds:

$$Mx = 0.$$

#### Optimization of kinetic proofreading performance

The objective of kinetic proofreading is to effectively discriminate between cognate and noncognate substrates, ensuring that the majority of the product consists of the correct molecule.

Conceptually, the system can be viewed as a discriminative pipeline, where the concentrations of cognate and noncognate substrates serve as inputs, so for simplicity, we assume that $x_R = x_W$ for the performance evaluation.

Therefore, the key parameters that can be varied in the system are:

$$k_{\text{on}},\ k_{\text{transit}},\ k_{\text{cat}},\ k_{\text{off}}^{\text{ER}},\ k_{\text{off}}^{\text{EW}}$$

The performance of this system is assessed in terms of activity $c$ and fidelity $\eta$. To optimize the system's performance, we seek to vary these parameters. The corresponding optimization problem is formulated as follows:

$$\begin{aligned} \min_{k} \quad & \eta, c \\ \text{s.t.} \quad & Mx = 0, \\ & \log \eta = \log x_{ER} - \log x_{EW}, \\ & \log c = \log x_{ER} \end{aligned} \tag{1}$$

Note that $n$ is treated as a fixed parameter, rather than a variable in this optimization problem.

This is because if $n$ are allowed to vary freely, the problem would become under-constrained, and potentially, leading to arbitrarily high performance. Specifically, as $n$ increases, if $k_{\text{transit}}$ were also allowed to increase without restriction, there would be no penalty on activity. However, freely increasing $n$ and $k_{\text{transit}}$ is not considered to be feasible within the current framework of kinetic proofreading.

The optimization problem has two objectives. Our goal is to determine the Pareto-optimal frontier, which represents the trade-off between the two objectives. To explore this trade-off, we constrain one objective while optimizing the other, yielding the following optimization problem, with the activity $c$ constrained.

$$\begin{aligned} \min_{k} \quad & \eta \\ \text{s.t.} \quad & Mx = 0, \\ & \eta = x_{ER}/x_{EW}, \\ & c = x_{ER} \end{aligned} \tag{2}$$

#### Solution of fidelity-activity trade-off in $n = 1$ and separable case

Here we provide a detailed description of the process for solving the optimization problem in the case where $n = 1$ and the system is separable. The term "Separable" refers to the assumption that we assume the competition for the enzyme molecule $E$ between the cognate and noncognate substrates is weak. Under this assumption, we can treat the kinetic proofreading processes for each substrate independently.

In this case, the state vector and the corresponding transition matrices are as follows:

$$x = \begin{bmatrix} x_E \\ x_{ES,0} \\ x_{ES,1} \end{bmatrix}, \; M_S = \begin{pmatrix} -k_{\text{on}} & k_{\text{off}}^{\text{ES}} & k_{\text{off}}^{\text{ES}} + k_{\text{cat}} \\ k_{\text{on}} & -(k_{\text{off}}^{\text{ES}} + k_{\text{transit}}) & 0 \\ 0 & k_{\text{transit}} & -(k_{\text{off}}^{\text{ES}} + k_{\text{cat}}) \end{pmatrix}$$

The steady state solution is contained in the unique eigenvector of the null space of $M_S$. From this, we obtain the steady state solution up to a multiplicative constant $c_0$, which is the total enzyme concentration $E_{tot} = c_0$.

Thus,

$$[\text{ER}_1] = c_0 \cdot \frac{k_{\text{on}} k_{\text{transit}}}{(k_{\text{off}}^{\text{ER}} + k_{\text{on}})(k_{\text{off}}^{\text{ER}} + k_{\text{transit}}) + k_{\text{cat}}(k_{\text{off}}^{\text{ER}} + k_{\text{on}} + k_{\text{transit}})}. \tag{3}$$

$$[\text{EW}_1] = c_0 \cdot \frac{k_{\text{on}} k_{\text{transit}}}{(k_{\text{off}}^{\text{EW}} + k_{\text{on}})(k_{\text{off}}^{\text{EW}} + k_{\text{transit}}) + k_{\text{cat}}(k_{\text{off}}^{\text{EW}} + k_{\text{on}} + k_{\text{transit}})}. \tag{4}$$

From this, the fidelity and activities are:

$$\eta_{n=1} = \frac{[\text{ER}_1]}{[\text{EW}_1]} = \frac{(k_{\text{off}}^{\text{EW}} + k_{\text{on}})(k_{\text{off}}^{\text{EW}} + k_{\text{transit}}) + k_{\text{cat}}(k_{\text{off}}^{\text{EW}} + k_{\text{on}} + k_{\text{transit}})}{(k_{\text{off}}^{\text{ER}} + k_{\text{on}})(k_{\text{off}}^{\text{ER}} + k_{\text{transit}}) + k_{\text{cat}}(k_{\text{off}}^{\text{ER}} + k_{\text{on}} + k_{\text{transit}})}. \tag{5}$$

$$c'_{n=1} = \frac{1}{c_0}[\text{ER}_1] = \frac{k_{\text{on}} k_{\text{transit}}}{(k_{\text{off}}^{\text{ER}} + k_{\text{on}})(k_{\text{off}}^{\text{ER}} + k_{\text{transit}}) + k_{\text{cat}}(k_{\text{off}}^{\text{ER}} + k_{\text{on}} + k_{\text{transit}})}. \tag{6}$$

We can substitute the explicit solutions obtained in this case to eliminate the $x_{ER}, x_{EW}$ variables in the optimization problem. The optimization problem then becomes:

$$\begin{aligned} \min_{k} \quad & \eta_{n=1}(k) \\ \text{s.t.} \quad & c_{n=1} = c(k) \end{aligned} \tag{7}$$

Here, we are subject to the constraint $c'_{n=1} = c$. We rewrite $\eta_{n=1}$ and $c'_{n=1}$ as $\eta_1$ and $c'_1$, respectively.

To determine the optimal value, we derive the Karush-Kuhn-Tucker conditions, which are necessary conditions for optimality.

The Lagrangian is,

$$\mathcal{L} = \eta_1 + \lambda \left( c'_1 - c \right), \tag{8}$$

We compute the partial derivatives $\frac{\partial \mathcal{L}}{\partial x} = 0$ for $x = (k_{\text{on}}, k_{\text{transit}}, k_{\text{cat}}, k_{\text{off}}^{\text{ER}}, k_{\text{off}}^{\text{EW}}, \lambda)$:

$$\frac{\partial \mathcal{L}}{\partial x} = \frac{\partial \eta_1}{\partial x} + \lambda \frac{\partial c'_1}{\partial x}, \quad \text{with respect to } x = \left(k_{\text{on}}, k_{\text{transit}}, k_{\text{cat}}, k_{\text{off}}^{\text{ER}}, k_{\text{off}}^{\text{EW}}, \lambda\right) \tag{9}$$

Define:

$$D_R = (k_{\text{off}}^{\text{ER}} + k_{\text{on}})(k_{\text{off}}^{\text{ER}} + k_{\text{transit}}) + k_{\text{cat}}(k_{\text{off}}^{\text{ER}} + k_{\text{on}} + k_{\text{transit}}), \tag{10}$$

$$D_W = (k_{\text{off}}^{\text{EW}} + k_{\text{on}})(k_{\text{off}}^{\text{EW}} + k_{\text{transit}}) + k_{\text{cat}}(k_{\text{off}}^{\text{EW}} + k_{\text{on}} + k_{\text{transit}}). \tag{11}$$

Then:

$$\eta_1 = \frac{D_W}{D_R} \tag{12}$$

As an example, take the derivative with respect to $k_{on}$,

$$\frac{\partial \mathcal{L}}{\partial k_{\text{on}}} = \frac{(k_{\text{off}}^{\text{ER}} + k_{\text{transit}} + k_{\text{cat}})D_W - D_R(k_{\text{off}}^{\text{EW}} + k_{\text{transit}} + k_{\text{cat}})}{D_W^2} + \lambda \frac{k_{\text{transit}} D_R - k_{\text{on}} k_{\text{transit}}(k_{\text{off}}^{\text{ER}} + k_{\text{transit}} + k_{\text{cat}})}{D_R^2} = 0. \tag{13}$$

The derivatives for other variables are computed similarly for $k_{\text{off}}^{\text{ER}}$, $k_{\text{off}}^{\text{EW}}$, $k_{\text{cat}}$, $k_{\text{transit}}$, $\lambda$. Each partial derivative $\frac{\partial \mathcal{L}}{\partial x} = 0$ forms a system of equations to be solved for the optimality condition.

Combining the partial derivatives with respect to $k_{\text{on}}$ and $k_{\text{transit}}$, we derive the following relationship:

$$\frac{\partial \mathcal{L}}{\partial k_{\text{on}}} - \frac{\partial \mathcal{L}}{\partial k_{\text{transit}}} = (k_{\text{transit}} - k_{\text{on}}) \left[ \frac{D_W - D_R}{D_W^2} + \lambda \frac{D_R - k_{\text{on}} k_{\text{transit}}}{D_R^2} \right]. \tag{14}$$

This equation, derived from the optimality conditions, underscores the relationship between $k_{\text{on}}$ and $k_{\text{transit}}$ in maximizing fidelity ($\eta_1$)while satisfying the constraint $c_1' - c = 0$.

From this, we can conclude that one of the optimality conditions is:

$$k_{\text{transit}} = k_{\text{on}}. \tag{15}$$

Under the conditions $k_{\text{transit}} = k_{\text{on}}$ and $k_{\text{off}}^{\text{ER}} \ll k_{\text{off}}^{\text{EW}}$, the normalized fidelity $\eta' = \frac{\eta}{\frac{k_{\text{off}}^{\text{EW}}}{k_{\text{off}}^{\text{ER}}}}$ simplifies to:

$$\eta_1' \approx \frac{(k_{\text{off}}^{\text{ER}})^2}{(k_{\text{off}}^{\text{ER}} + k_{\text{on}})^2 + k_{\text{cat}}(k_{\text{off}}^{\text{ER}} + 2k_{\text{on}})}. \tag{16}$$

The term $c_1'$ is calculated as:

$$c_1' = \frac{k_{\text{on}}^2}{(k_{\text{off}}^{\text{ER}} + k_{\text{on}})^2 + k_{\text{cat}}(k_{\text{off}}^{\text{ER}} + 2k_{\text{on}})}. \tag{17}$$

Thus, the expression for $\eta_1' + c_1'$ is given by:

$$\eta_1' + c_1' = \frac{(k_{\text{off}}^{\text{ER}})^2 + k_{\text{on}}^2}{(k_{\text{off}}^{\text{ER}} + k_{\text{on}})^2 + k_{\text{cat}}(k_{\text{off}}^{\text{ER}} + 2k_{\text{on}})}. \tag{18}$$

Under the condition where $k_{\text{cat}} \to 0$, we derive the upper boundary (see Figure 5C) for the activity-fidelity trade-off:

$$(\eta_1')^{1/2} + (c_1')^{1/2} \to 1. \tag{19}$$

#### Generalized trade-off between activity and fidelity for any n

We analyze the reactions for $n > 1$. For example, when $n = 2$, the reaction matrix is:

$$M_2 = \begin{pmatrix} -k_{\text{on}} & k_{\text{off}}^{\text{ER}} & k_{\text{off}}^{\text{ER}} & k_{\text{off}}^{\text{ER}} + k_{\text{cat}} \\ k_{\text{on}} & -(k_{\text{off}}^{\text{ER}} + k_{\text{transit}}) & 0 & 0 \\ 0 & k_{\text{transit}} & -(k_{\text{off}}^{\text{ER}} + k_{\text{transit}}) & 0 \\ 0 & 0 & k_{\text{transit}} & -(k_{\text{off}}^{\text{ER}} + k_{\text{cat}}) \end{pmatrix} \tag{20}$$

For $n = 3$:

$$M_3 = \begin{pmatrix} -k_{\text{on}} & k_{\text{off}}^{\text{ER}} & k_{\text{off}}^{\text{ER}} & k_{\text{off}}^{\text{ER}} & k_{\text{off}}^{\text{ER}} + k_{\text{cat}} \\ k_{\text{on}} & -(k_{\text{off}}^{\text{ER}} + k_{\text{transit}}) & 0 & 0 & 0 \\ 0 & k_{\text{transit}} & -(k_{\text{off}}^{\text{ER}} + k_{\text{transit}}) & 0 & 0 \\ 0 & 0 & k_{\text{transit}} & -(k_{\text{off}}^{\text{ER}} + k_{\text{transit}}) & 0 \\ 0 & 0 & 0 & k_{\text{transit}} & -(k_{\text{off}}^{\text{ER}} + k_{\text{cat}}) \end{pmatrix} \tag{21}$$

For $n > 1$, steady-state solutions and fidelity are derived analogously to the $n = 1$ case, with increased matrix dimensions reflecting additional intermediate states. By applying the method described for $n = 1$, we determine the concentration and fidelity for any $n$ to be:

$$c'_n = \frac{k_{\text{on}}(k_{\text{transit}})^n}{(k_{\text{off}}^{\text{ER}} + k_{\text{on}})(k_{\text{off}}^{\text{ER}} + k_{\text{transit}})^n + k_{\text{cat}} \cdot f_n(k_{\text{off}}^{\text{ER}}, k_{\text{on}}, k_{\text{transit}})} \tag{22}$$

$$\eta_n = \frac{(k_{\text{off}}^{\text{EW}} + k_{\text{on}})(k_{\text{off}}^{\text{EW}} + k_{\text{transit}})^n + k_{\text{cat}} \cdot f_n(k_{\text{off}}^{\text{EW}}, k_{\text{on}}, k_{\text{transit}})}{(k_{\text{off}}^{\text{ER}} + k_{\text{on}})(k_{\text{off}}^{\text{ER}} + k_{\text{transit}})^n + k_{\text{cat}} \cdot f_n(k_{\text{off}}^{\text{ER}}, k_{\text{on}}, k_{\text{transit}})} \tag{23}$$

Here, the $f_n$ refers to:

$$f_n(k_{\text{off}}, k_{\text{on}}, k_{\text{transit}}) = (k_{\text{off}} + k_{\text{transit}})^n + k_{\text{on}} \cdot \sum_{i=0}^{n-1} \binom{n-1}{i} (i+1) k_{\text{off}}^{n-1-i} (k_{\text{transit}})^i \tag{24}$$

Hence, for any $n$, similar Lagrange optimality conditions are derived to be $k_{\text{transit}} = k_{\text{on}}$, $k_{\text{off}}^{\text{ER}} \ll k_{\text{off}}^{\text{EW}}$, and $k_{\text{cat}} \to 0$. This regime yields the following relationship between activity and fidelity (shown in Figure 5C).

$$\eta'^{1/(n+1)}_n + c'^{1/(n+1)}_n = 1 \tag{25}$$

### Theoretical analysis of classic kinetic proofreading systems with competitive binding

#### Description of the system

We study the dynamics of kinetic proofreading reactions where the enzyme (E) binds to both cognate (R) and noncognate (W) substrates with the same association rate constant, $k_{\text{on}}$ and dissociation rate constants $k_{\text{off}}^{\text{ER}}$ or $k_{\text{off}}^{\text{EW}}$, respectively, forming the ER and EW complexes. **Document S1** considers the binding of cognate and noncognate complexes. Here, we consider them together, separately.

We start the analysis of proofreading reactions with one intermediate state (n = 1). The kinetic proofreading system involves the enzyme E interacting with both cognate (R) and noncognate (W) substrates. The system includes five states: free enzyme (E), enzyme-cognate complexes (ER$_0$, ER$_1$), and enzyme-noncognate complexes (EW$_0$, EW$_1$). Here, $k_{\text{transit}}$ denotes the transition rate for the kinetic proofreading steps. The final intermediate states ER$_1$ and EW$_1$ produce final products with the catalytic rate of $k_{\text{cat}}$.

The chemical reactions governing the system are as follows:

$$\text{E} + \text{R} \underset{k_{\text{off}}^{\text{ER}}}{\overset{k_{\text{on}}}{\rightleftharpoons}} \text{ER}_0,$$

$$\text{E} + \text{W} \underset{k_{\text{off}}^{\text{EW}}}{\overset{k_{\text{on}}}{\rightleftharpoons}} \text{EW}_0,$$

$$\text{ER}_0 \xrightarrow{k_{\text{transit}}} \text{ER}_1,$$

$$\text{EW}_0 \xrightarrow{k_{\text{transit}}} \text{EW}_1,$$

$$\text{ER}_\text{i} \xrightarrow{k_{\text{off}}} \text{E} + \text{R}\ (i = 0, 1),$$

$$\text{EW}_\text{i} \xrightarrow{k_{\text{off}}} \text{E} + \text{W}\ (i = 0, 1),$$

$$\text{ER}_1 \xrightarrow{k_{\text{cat}}} \text{E} + \text{P}_\text{R},$$

$$\text{EW}_1 \xrightarrow{k_{\text{cat}}} \text{E} + \text{P}_\text{W}.$$

We introduce the vectors of molecular concentrations $x$ as:

$$x = \begin{bmatrix} x_{\text{E}} \\ x_{\text{ER}_0} \\ x_{\text{ER}_1} \\ x_{\text{EW}_0} \\ x_{\text{EW}_1} \end{bmatrix} \tag{1}$$

The dynamics of this system are described by:

$$\frac{dx}{dt} = Mx \tag{2}$$

where M is transition rate matrix.

#### Steady state analysis ($n = 1$)

The transition rate matrix M is:

$$M = \begin{pmatrix} -2k_{\text{on}} & k_{\text{off}}^{\text{ER}} & k_{\text{off}}^{\text{ER}} + k_{\text{cat}} & k_{\text{off}}^{\text{EW}} & k_{\text{off}}^{\text{EW}} + k_{\text{cat}} \\ k_{\text{on}} & -(k_{\text{off}}^{\text{ER}} + k_{\text{transit}}) & 0 & 0 & 0 \\ 0 & k_{\text{transit}} & -(k_{\text{off}}^{\text{ER}} + k_{\text{cat}}) & 0 & 0 \\ k_{\text{on}} & 0 & 0 & -(k_{\text{off}}^{\text{EW}} + k_{\text{transit}}) & 0 \\ 0 & 0 & 0 & k_{\text{transit}} & -(k_{\text{off}}^{\text{EW}} + k_{\text{cat}}) \end{pmatrix} \tag{3}$$

To investigate the performance of kinetic proofreading, we focus on the steady-state behavior of the kinetic proofreading system. At steady state, the following condition holds:

$$Mx = 0.$$

Solving the steady-state equations and applying the enzyme conservation constraint:

$$[\text{E}] + [\text{ER}_0] + [\text{ER}_1] + [\text{EW}_0] + [\text{EW}_1] = c_0 \tag{4}$$

Define:

$$D_R = (k_{\text{off}}^{\text{ER}} + k_{\text{transit}})(k_{\text{off}}^{\text{ER}} + k_{\text{cat}}) \tag{5}$$

$$D_W = (k_{\text{off}}^{\text{EW}} + k_{\text{transit}})(k_{\text{off}}^{\text{EW}} + k_{\text{cat}}) \tag{6}$$

The steady-state concentrations of the final complexes are:

$$[\text{ER}_1] = c_0 \cdot \frac{k_{\text{on}} k_{\text{transit}}}{D_R + k_{\text{on}}(k_{\text{off}}^{\text{ER}} + k_{\text{transit}} + k_{\text{cat}}) + \frac{D_R}{D_W} k_{\text{on}}(k_{\text{off}}^{\text{EW}} + k_{\text{transit}} + k_{\text{cat}})} \tag{7}$$

$$[\text{EW}_1] = c_0 \cdot \frac{k_{\text{on}} k_{\text{transit}}}{D_W + k_{\text{on}}(k_{\text{off}}^{\text{EW}} + k_{\text{transit}} + k_{\text{cat}}) + \frac{D_W}{D_R} k_{\text{on}}(k_{\text{off}}^{\text{ER}} + k_{\text{transit}} + k_{\text{cat}})} \tag{8}$$

#### Circuit fidelity and activity ($n = 1$)

Since the total concentration of E is a constant (Eq (4)) , we can use the normalized concentration of $\text{ER}_n$ in the following derivation, where $c' c_0 = c_n$. According to Eq (7) and Eq (8) , the normalized activity here is:

$$c'_1 = \frac{[\text{ER}_1]}{c_0} = \frac{k_{\text{on}} k_{\text{transit}}}{D_R + k_{\text{on}}(k_{\text{off}}^{\text{ER}} + k_{\text{transit}} + k_{\text{cat}}) + \frac{D_R}{D_W} k_{\text{on}}(k_{\text{off}}^{\text{EW}} + k_{\text{transit}} + k_{\text{cat}})} \tag{9}$$

The fidelity is:

$$\eta_1 = \frac{[\text{ER}_1]}{[\text{EW}_1]} = \frac{D_W + k_{\text{on}}(k_{\text{off}}^{\text{EW}} + k_{\text{transit}} + k_{\text{cat}}) + \frac{D_W}{D_R} k_{\text{on}}(k_{\text{off}}^{\text{ER}} + k_{\text{transit}} + k_{\text{cat}})}{D_R + k_{\text{on}}(k_{\text{off}}^{\text{ER}} + k_{\text{transit}} + k_{\text{cat}}}) + \frac{D_R}{D_W} k_{\text{on}}(k_{\text{off}}^{\text{EW}} + k_{\text{transit}} + k_{\text{cat}}) \tag{10}$$

$$= \frac{(k_{\text{cat}} + k_{\text{off}}^{\text{EW}})((k_{\text{transit}} + k_{\text{off}}^{\text{EW}}))}{(k_{\text{cat}} + k_{\text{off}}^{\text{ER}})(k_{\text{transit}} + k_{\text{off}}^{\text{ER}})} \tag{11}$$

These results show the activity-fidelity trade-off in a kinetic proofreading system (see Figure S9), with the competition between cognate and noncognate substrates for the enzyme.
